# Friends with malefit. The effects of keeping dogs and cats, sustaining animal-related injuries and *Toxoplasma* infection on health and quality of life

**DOI:** 10.1101/742734

**Authors:** Jaroslav Flegr, Marek Preiss

## Abstract

Many studies show that keeping cats and dogs has a positive impact on humans’ physical and mental health and quality of life. The existence of this “pet phenomenon” is now widely discussed because other studies performed recently have demonstrated a negative impact of owning pets or no impact at all. The main problem of many studies was the autoselection – participants were informed about the aims of the study during recruitment and later likely described their health and wellbeing according to their personal beliefs and wishes, not according to their real status. To avoid this source of bias, we did not mention pets during participant recruitment and hid the pet-related questions among many hundreds of questions in an 80-minute Internet questionnaire. Results of our study performed on a sample of on 10,858 subjects showed that liking cats and dogs has a weak positive association with quality of life. However, keeping pets, especially cats, and even more being injured by pets, were strongly negatively associated with many facets of quality of life. Our data also confirmed that infection by the cat parasite *Toxoplasma* had a very strong negative effect on quality of life, especially on mental health. However, the infection was not responsible for the observed negative effects of keeping pets, as these effects were much stronger in 1,527 *Toxoplasma*-free subjects than in the whole population. Any cross-sectional study cannot discriminate between a cause and an effect. However, because of the large and still growing popularity of keeping pets, the existence and nature of the reverse pet phenomenon deserve the outmost attention.

## Introduction

According to 2016 estimates, about 61 million dogs and 66 million cats are currently kept in the EU and this number is likely still growing [1]. Keeping dogs and cats is believed to have many positive effects on the wellbeing and health of humans. Since the eighties, many papers have been published showing that contact with pets has positive effects on individuals facing various life stressors [2](Allen 2001) as well as on normal people suffering from loneliness [3] and depression [4–6]. In the general population, keeping pets, and especially dogs, was shown to have positive effects on the subjects’ community connectedness, and on their mental health and wellbeing [7–9]. It was also shown that keeping dogs, but often not cats [10], increased the chance for survival after heart diseases including myocardial infarction [10] and even decreased the total mortality rate in various populations [11–14]. Simply engaging with animals, e.g. stroking animals or just thinking about pets, was shown to reduce blood pressure and stress [15] and to provide relief from social rejection [16]. Pet owners were shown to spend a smaller fraction of their income on healthcare than non-owners [17].

However, in the past twenty years, more and more studies were performed that failed to reproduce the results of these older studies. They either observed no effects of keeping pets on human wellbeing and health [18], or actually observed negative effects of keeping dogs, and even more of keeping cats [19], on health, survival of patients and members of general population and their wellbeing [13, 20–30]. The question of the existence of positive or negative effects of keeping pets is still open; however, more and more authors have reached the conclusion that many of the older results may be strongly biased by autoselection of participants of studies and selective reporting of only positive results, or *a priori* expected results of studies [17]. Also, even many highly cited studies were performed on rather small or imbalanced samples of participants and most of them reported only the effects of dogs, despite the fact that data on the effects of cats were probably also studied [8, 9, 31, 32]. It is sometimes stated that no differences in economic situation exists between the pet keepers and non keepers [33]; however, this is probably not true as the cost of keeping a dog and a cat (which usually lives longer) is estimated at $8,000 and $10,000, respectively [17]. The economic situation of subjects is correlated with their health and probably also their quality of life. It is also found that old people keep pets less often then middle age people, often because, for technical and economic reasons, they cannot meet the needs of their pets [27, 34]. The age of subjects is usually controlled in statistical tests. However, biological age (and health) as opposed to calendar age is probably responsible for the observed negative association between keeping pets and age of participants of study. This means, paradoxically, that by controlling for calendar age we can introduce further bias into the data – only people with aberrantly low biological age are able to keep pets in late senior age. The most common nontrivial problem of earlier studies was that the participants in the studies were informed in advance that the effects of pets (or dogs and cats, or animals) on quality of life will be the subject of the study. It is highly probable that people who keep pets also love them and believe that pets make their life better (otherwise they would get rid of them) and therefore answer the questions in agreement with their *a priori* beliefs, regardless of the real effects of pets on their lives. This is in agreement with a common observation that, when directly asked, people often report a positive effect of their pets on wellbeing while the results of detailed questionnaires suggest an absence of such effects [30].

To avoid this problem we ran a large Internet study that was advertised without mentioning pets, cats, dogs or animals. The questions related to dogs and cats were buried within many hundreds of other unrelated questions in different parts of the questionnaire than the health- and wellbeing-related questions. This made it rather improbable that dog- and cat-keepers/lovers would answer the quality of life- and health-related questions in accordance with their personal opinions on the effects of pets on quality of human life. In a study performed on 10,858 subjects, we searched not only for the effects of keeping pets but also for the effects of loving cats and sustaining animal-related injuries. It is often suggested that latent infection with the cat parasite *Toxoplasma* (whose oocysts can be transmitted by both cat and dogs with soil on paws) is responsible for impaired mental and physical health of people. To test this hypothesis, we also studied the association of pet-related variables and health and wellbeing in a subpopulation of 343 men and 1,184 women who had been tested negatively for toxoplasmosis.

## Materials and methods

### Subjects

The internet questionnaire was distributed as a Qualtrics survey. Subjects were invited to participate in the study using a Facebook-based snowball method [35]. Potential volunteers, mostly members of the “Lab bunnies” community, an 18,000-member group of Czech and Slovak nationals willing to take part in evolutionary psychology experiments, and their Facebook friends, were invited (using about 10 different posts on the Lab bunnies timeline) to participate in anonymous study about “mystical thinking, superstitions, prejudices, religion and the relation between various environmental factors and health and wellbeing.” The electronic questionnaire was also promoted in various electronic and printed media and TV. Keeping cats and dogs (or “pets” or “animals”) was not explicitly mentioned in the information provided to potential participants. Responders were not paid for their participation in the study; however, after finishing the 80-minute questionnaire, they were provided information about the results of related studies and their own results of several tests that were part of the questionnaire. At the first screen of the survey, the participants were given the following information and were asked to provide their informed consent to participate in the study by pressing a special button: “The study is anonymous and obtained data will be used exclusively for scientific purposes. Your cooperation in the project is voluntary and you can terminate it at any time by closing this web page. You can also skip any uncomfortable questions; however, complete data is most valuable. If you agree to participate in the research press the “Next” button”. Only the subjects who provided their informed consent were allowed to participate in the study. Some pages of the questionnaire contained the Facebook share button. These buttons were pressed by more than 1,600 participants, which resulted in obtaining data from 12,600 responders in total between 27^th^ May 2016 and 29^th^ June 2018. All methods were performed in accordance with the relevant guidelines and regulations. The project, including the method of obtaining electronic informed consent to participate in this anonymous study from all participants, was approved by the IRB of the Faculty of Science, Charles University (Komise pro práci s lidmi a lidským materiálem Přírodovědecké Fakulty Univerzity Karlovy) - No. 2015/07.

### Questionnaires

The electronic survey consisted of several parts that concerned various unrelated projects on evolutionary psychology and psychiatry. In the present study, we inspected and analyzed only responses to the questions concerning health, wellbeing, biological fitness and keeping dogs and cats. The responders were asked about their *sex, age, education* (ordinal scale 1-8: 1-elementary, 8-PhD or MD), *body height*, *body weight*, and the size of the communities where they currently live (ordinal variable *urbanization*: 0-less than 1000 inhabitants, 1-1-5 thousand inhabitants, 2: 5-50 thousand inhabitants, 3: 50-100 thousand inhabitants, 4: 100-500 thousand inhabitants, 5: more than 500 thousand inhabitants). The anamnestic part of the questionnaire contained several questions about the intensity and nature of the persońs contact with dogs and cats and about sustained animal-related injuries. Subjects were asked to rate how much they *like dogs (cats)* using the scale 0-100 (the *preference of dogs to cats* was calculated as the difference between liking dogs and liking cats), and then rate the intensity of their past and current contacts with dogs and cats using a 8-points pseudo-scale: (0: our family never kept a dog (cat), 1-we kept a dog (cat) in the past but only for a short time, 2-we kept a dog (cat) only in the past but for several years, 4-we have one dog (cat), 5-we have two dogs (cats), 6-we have three dogs (cats), 7-we have more than three dogs (cats)). Based on their responses, we calculated three variables describing the intensity of contacts with dogs and three with cats: *Ever keeping dog (cat)* (binary: codes 0 vs 1-7), *Now keeping a dog (cat)* (binary: codes 0, 1, 2 vs 3-7), *Number of dogs (cats)* in house (ordinal: 3-7). Next, the responders were asked to rate the intensity of sustained animal-related injuries (three variables: *biting by a dog, biting by a cat, and scratching by a cat*) using the following scales: 0-never, 1-only while playing, 2-only as a warning, 3-yes, minor injury (only skin cut), 4-yes, moderate injury (bleeding), 5-yes, I had to seek medical treatment, 6-I was seriously injured by several dogs (cats). For the purposes of future analyses, we merged the infrequently used category 6 with category 5. To see, whether toxoplasmosis could play a role on studied associations, we also asked the responders whether they had been laboratory tested for this infection, what was the result of this test (binary variable *toxoplasmosis,* the third response “I do not know, I am not sure” was *a priory* checked), and what was the purpose of this testing (categorical: research in our lab, health reasons, pregnancy, other). As a benchmark for the relative importance of associations of the animals-related variables with the health, wellbeing, and biological fitness, we looked for the associations of health, wellbeing, and fitness with four unrelated but well-known risk factors: *body mass index* (BMI) calculated from body height and body weight, frequency of *smoking*, i.e., how many cigarettes they smoke a day (ordinal scale: 0-0, 1-0-0.1, 2-0.1-1, 3-1.1-3, 4-3.1-10, 5-11-20, 6-21-40, 7-more than 40), frequency of *consuming alcohol* “not to be allowed to drive a car for a while for this reason” (ordinal scale: 0-never, 1-maximally 1× a months, 2-maximally 2× a months, 3-maximally 4× a months, 4-maximally 2× a week, 5-every second day, 6-every day, 7-nearly all the time), and frequency of *consuming illegal drugs* “not to be allowed to drive a car for a while for this reason” (ordinal scale: 0-never, 1-maximally 1× a months, 2-maximally 2× a months, 3-maximally 4× a months, 4-maximally 2× a week, 5-every second day, 6-every day, 7-nearly all the time).

In another part of the questionnaire we collected information concerning the following outcome variables from the responders: How they rate their physical health status in comparison with other people of the same age (subjective *physical health problems*: six points scale, anchored with 0-definitively better status, 5-definitively worse status), How they rate their mental health status in comparison with other people of the same age (*subjective mental health problems*: the same scale), how they rate their *family situation*, e.g. the quality of an emotional support they can receive, (0-poor, 5-excellent), how they rate the *economic situation* of their family (0-poor, 5-excellent). To obtain more objective and concrete information on the health status of responders, we also asked them the following questions: how many kinds of *drugs prescribed* by a medical doctor they were taking currently, how many kinds of *drugs non-prescribed* by a doctor they were taking currently (“how many different herbs, food supplements, multivitamins, superfoods etc. do you currently take per day”), how many times they visited their primary care doctor in past 365 days (“not for prevention”), how many times they used *antibiotics* in the past 365 days, and how many different *medical specialists* they visited (not for prevention) in the past 5 years. The *physical health problems score* was calculated as a mean of Z-scores of the last five variables. The responders were also requested to rate how much they suffer with *anxieties, phobias, depression, mania, obsessions, auditory hallucinations, visual hallucinations,* and *headaches* using a 0-100 scale. We also counted *number of diagnosed* and *number of undiagnosed mental health disorders* of responders and of their partners, all checked on a list of 25 mental health disorders and epilepsy. The *mental health problems score* was calculated as a mean of Z-scores of the last 10 variables (not the numbers of diagnosed and undiagnosed disorders of the responder’s partner). Using a 0-100 scale they also answered the questions - How intensely they are *sexually attracted to men*, How intensely they are *sexually attracted to women*, (Z-score of the higher of these two responses was considered as the *intensity of sexual desire*). They were also asked with *how many men (women)* they had sex (“vaginal, oral or anal”). To respond to these two and the next two questions, the participants used a 0-9 ordinal scale (0-0, 1-1, 2-2, 3-3, 4-4, 5-5-6, 6-7-9, 7-10-19, 8-20 and more). The higher of these two responses was considered to reflect the number of *preferred sex sexual partners*. Similarly, the participants were asked with how many men (women) they exchanged in *French kissing* (the same ordinal scale) - the higher of these two responses was considered to reflect the number of *preferred sex French kissing*-*partners*. In the other part of the questionnaire, the responders were asked to estimate how many minutes daily they spend engaged in various activities. The list of 22 activities also contained “*any form of sex*, including consuming pornography”. *Score of sexual activity* was computed as mean of Z-scores of number of preferred sex sexual partners, preferred sex French kissing partners, and minutes spend engaged in any form of sex per day. As a proxy for direct and inclusive biological fitness we used number of biological *children* and number of *siblings,* respectively. For the assessment of quality of life of responders we used the Czech version of the standard 24-item WHOQOL-BREF (The World Health Organisation Quality of Life Assessment Instrument – the abbreviated version of the WHOQOL-100) [36]. This instrument had been translated into Czech and standardized to the Czech population [37]. It monitors general quality of life and its four specific domains: Physical health (activity of daily living, dependence on medical substances and aids, energy and fatigue, mobility, pain and discomfort, sleep and rest, work capacity), Psychological (bodily image and appearance, negative feelings, positive feelings, self-esteem, spirituality/religion/personal beliefs, thinking, learning, memory and concentration), Social relationships (personal relationships, social support, sexual activity), and Environment (financial resources, freedom, physical safety and security, health and social care: accessibility and quality, home environment, opportunity of acquiring new information and skills, participation in and opportunity for recreation/leisure activities, physical environment (pollution/noise/traffic/climate), transport).

### Statistical analyses

Before any analyses, records of all subjects who did not answer the pet-related questions and of about 2% of subjects who provided a suspicious combination of answers to other questions (too high/low body height, weight, age, an unrealistically high number of neuropsychiatric disorders, who answered all or nearly all questions by the same code, etc.) were filtered out.

The final set contained data from 10,858 subjects; however, not all of them responded to all questions concerning their health, quality of life, and keeping cats and dogs. The distribution of all relevant (semi-continuous and ordinal) variables was visually checked and then all secondary indices, see above, were computed. Statistical analysis was performed with the statistical package Statistica v.10.0. (descriptive statistics, t-tests, contingency tables, logistic regression). For computing the partial Kendall correlation we used R 3.3.1 and the package ppcor. The correction for multiple tests was done using Benjamini-Hochberg procedure with the false discovery rate pre-set to 0.20 [38]. The datasets generated during and/or analyzed during the current study are available in the figshare repository, https://figshare.com/s/58a34af3d3590dc2d398.

### Terminological note

Through the paper, the statistical relations between (formally) dependent and (formally) independent variables are called “effects,” despite the fact that the real causal relationship between these variables may be different or even non-existent (as pointed out in the Discussion).

## Results

### a) Descriptive statistics

The final sample consisted of 4,274 men (age 35.0, SD 12.6) and 6,584 women (age 34.7, SD 12.8), ns. Only 1131 (30.8%) men and 1,454 (24.8%) women never kept a dog (Chi^2=^42.1, d.f.=1, p<0.0001) and only 1,257 (34.4%) men and 1,680 (28.6%) women never kept a cat (Chi^2^=35.7, d.f.=1, p<0.0001). The people who ever kept a dog also had a higher probability of ever having kept a cat (Chi^2^=512.3, d.f.=1, p<0.0001). This result was confirmed by logistic regression with ever keeping a dog as the output binary variable and ever keeping a cat, sex, age, urbanization (size of place of living), and education as the predictors. Similar results showed the logistic analysis for ever keeping a cat as the output variable and ever keeping a dog, sex, age, size of urbanization, and education as the predictors (Tab. 1).

**Table 1.**
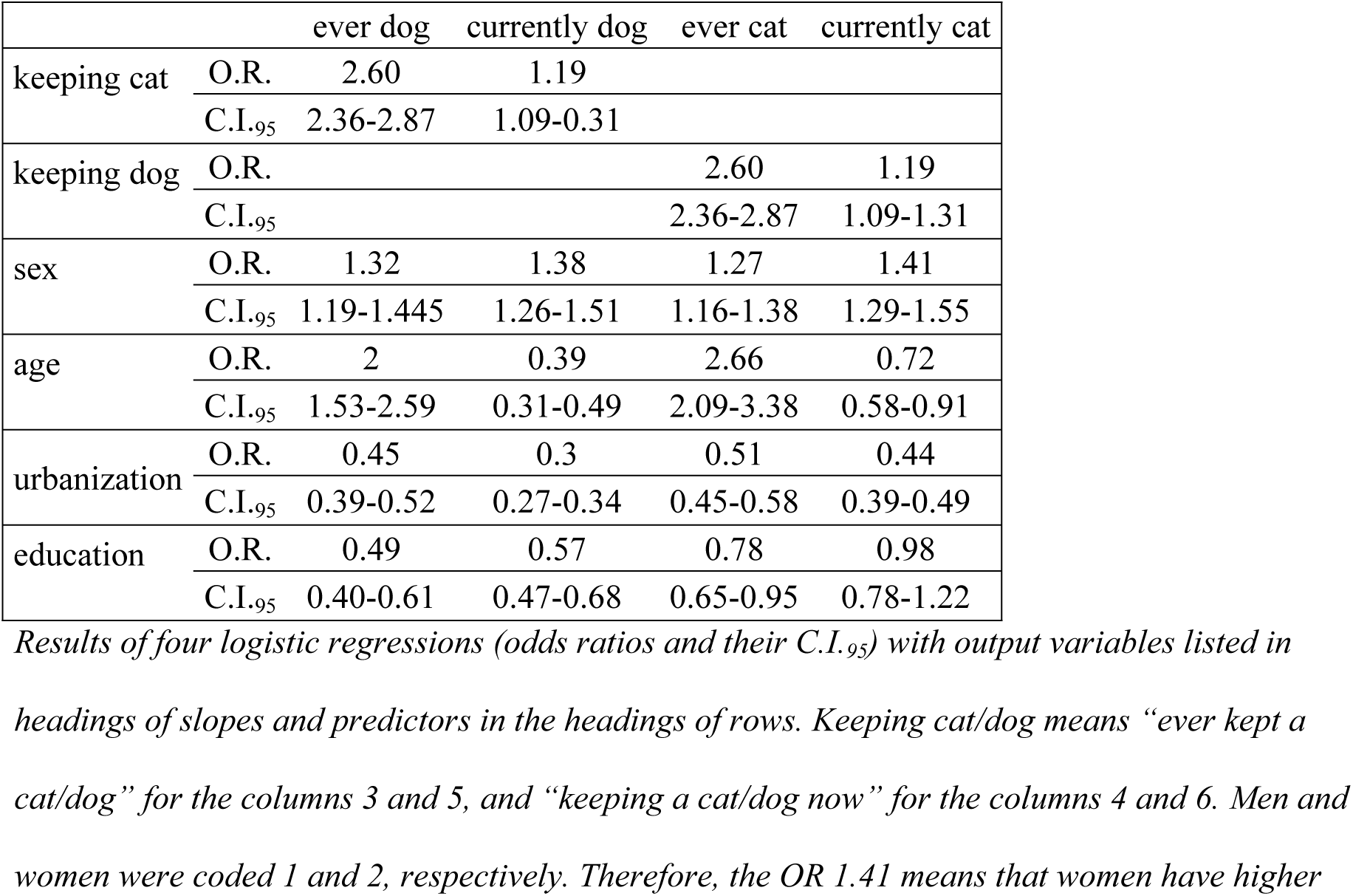
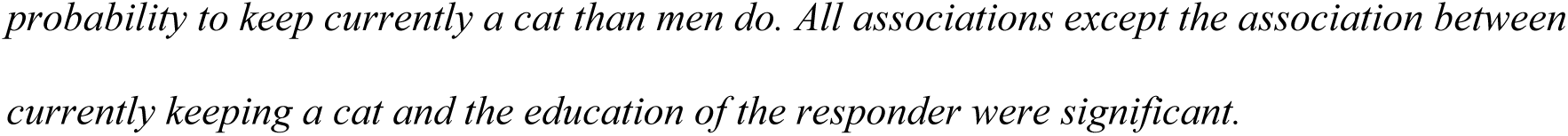
Predictors of keeping cats and dogs

|  |  | ever dog | currently dog | ever cat | currently cat |
| --- | --- | --- | --- | --- | --- |
| keeping cat | O.R. | 2.60 | 1.19 |  |  |
|  | C.I.<sub>.95</sub> | 2.36-2.87 | 1.09-0.31 |  |  |
| keeping dog | O.R. |  |  | 2.60 | 1.19 |
|  | C.I.<sub>.95</sub> |  |  | 2.36-2.87 | 1.09-1.31 |
| sex | O.R. | 1.32 | 1.38 | 1.27 | 1.41 |
|  | C.I.<sub>.95</sub> | 1.19-1.445 | 1.26-1.51 | 1.16-1.38 | 1.29-1.55 |
| age | O.R. | 2 | 0.39 | 2.66 | 0.72 |
|  | C.I.<sub>.95</sub> | 1.53-2.59 | 0.31-0.49 | 2.09-3.38 | 0.58-0.91 |
| urbanization | O.R. | 0.45 | 0.3 | 0.51 | 0.44 |
|  | C.I.<sub>.95</sub> | 0.39-0.52 | 0.27-0.34 | 0.45-0.58 | 0.39-0.49 |
| education | O.R. | 0.49 | 0.57 | 0.78 | 0.98 |
|  | C.I.<sub>.95</sub> | 0.40-0.61 | 0.47-0.68 | 0.65-0.95 | 0.78-1.22 |

*Results of four logistic regressions (odds ratios and their C.I.<sub>.95</sub>) with output variables listed in headings of slopes and predictors in the headings of rows. Keeping cat/dog means “ever kept a cat/dog” for the columns 3 and 5, and “keeping a cat/dog now” for the columns 4 and 6. Men and women were coded 1 and 2, respectively. Therefore, the OR 1.41 means that women have higher*
*probability to keep currently a cat than men do. All associations except the association between* *currently keeping a cat and the education of the responder were significant.*

At the time of the study, 1,147 (31.3%) men and 2,279 (38.8%) women kept a dog (Chi^2=^55.87, d.f.=1, p<0.0001) and 1,130 (30.9%) men and 2,303 (39.1%) women kept a cat (Chi^2^=66.68, d.f.=1, p<0.0001). Again, the people who were keeping a dog at the time of the study had a higher probability of keeping a cat at the time of the study (Chi^2^=56.70, d.f.=1, p<0.0001). This result was confirmed with logistic regressions, see the Table 1.

The more detailed structure of the analyzed population, namely the distribution of responses on particular questions of the questionnaire and arithmetic means of continuous dependent variables in men and women, are shown in Tables 2-3.

**Table 2.**
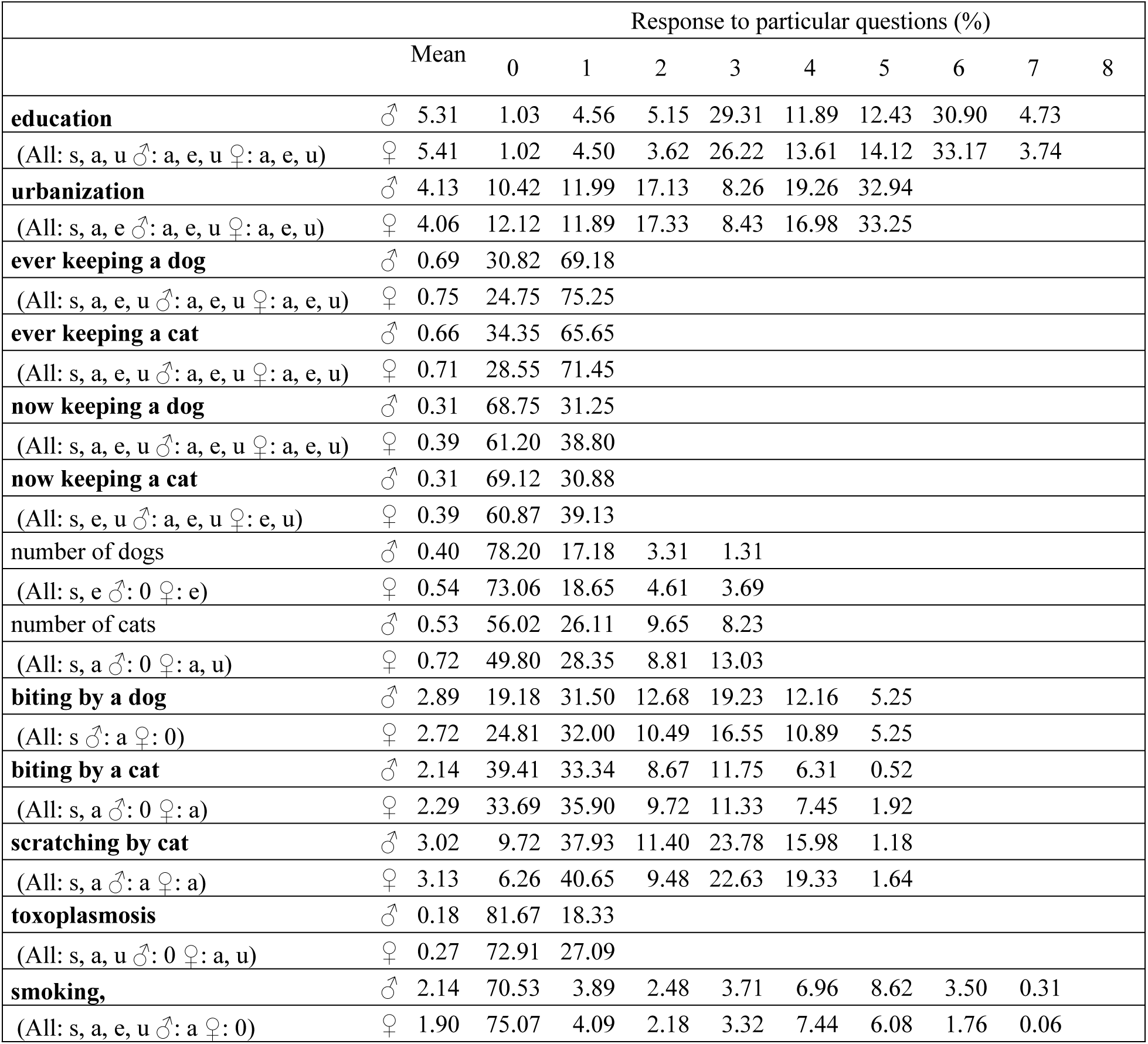
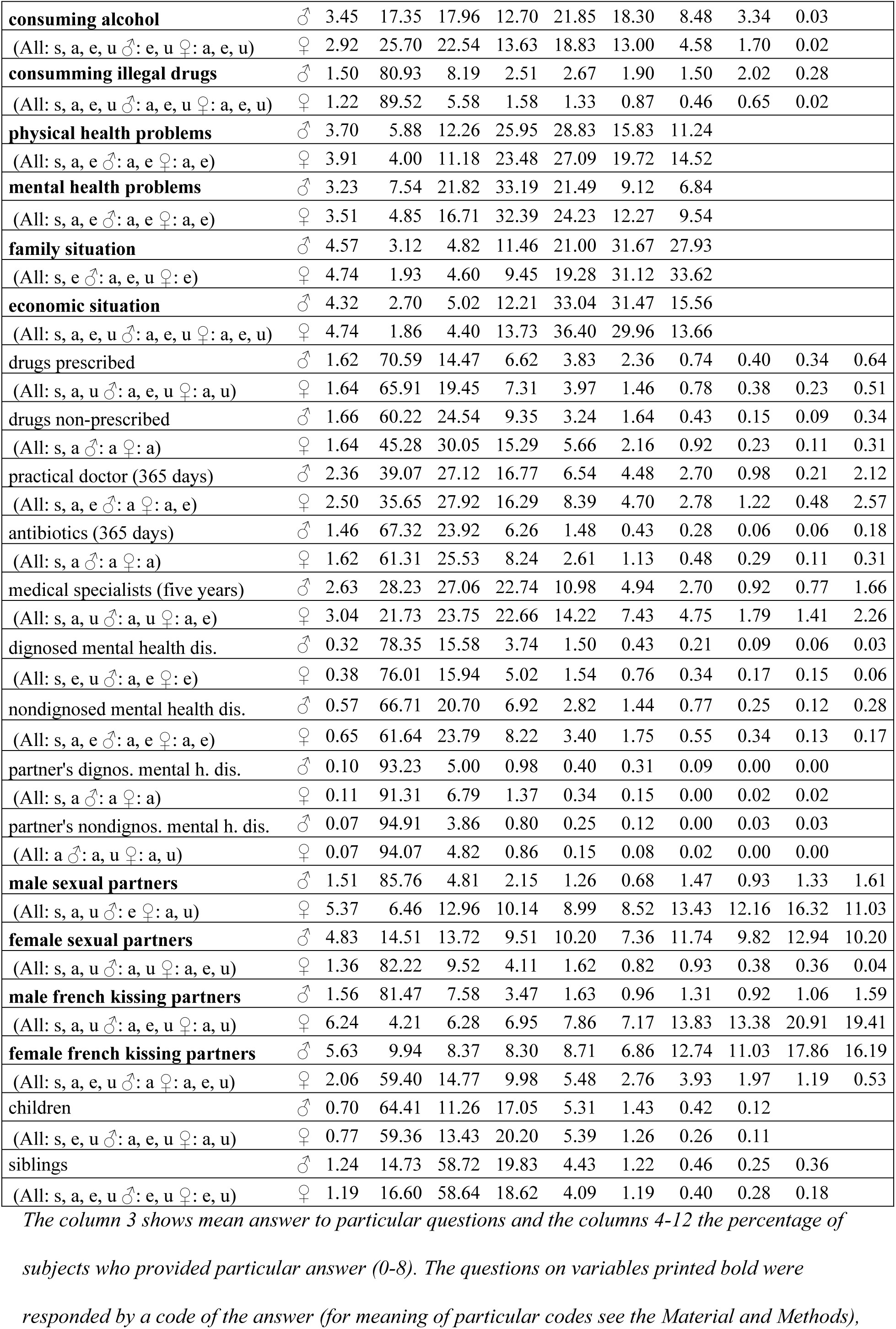
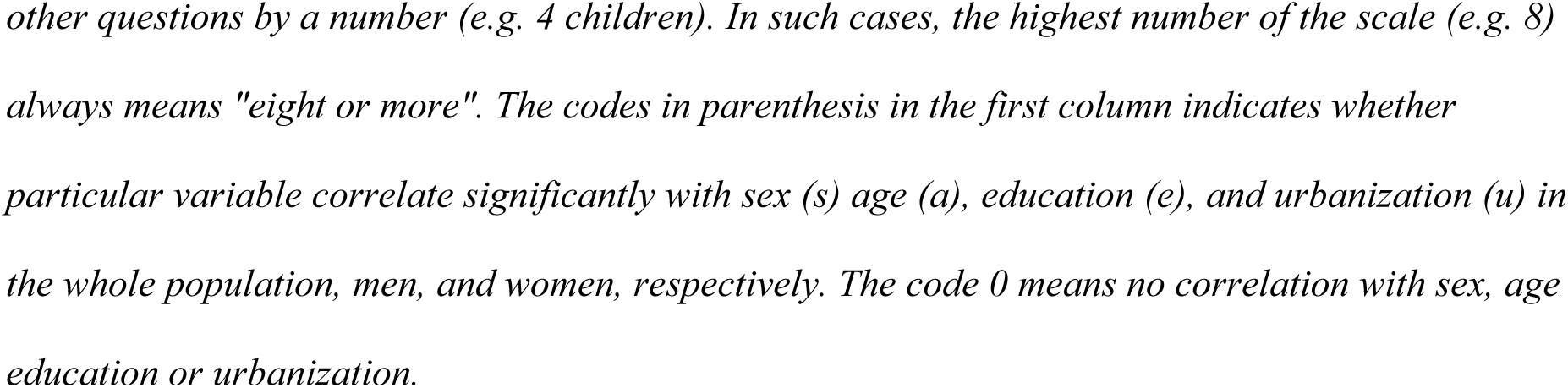
Distribution of responses on particular questions of the questionnaire – categorical and ordinal variables

**Table 3.** Continuous depending variables - difference between men and women.

|  | Mean Men | Mean Women | t-value | d.f. | p |
| --- | --- | --- | --- | --- | --- |
| Wellbeing (WHOQOL-BREF)<br>(All: a, e ♂: a ♀: a, e) | 97.267 | 97.091 | 0.536 | 7482 | 0.592 |
| Mental health problems score<br>(All: s, a, u ♂: a, e ♀: a, e) | -0.082 | 0.049 | -10.346 | 8583 | 0.000 |
| Physical health problems score<br>(All: s, a, e, u ♂: a ♀: a, e) | -0.089 | 0.057 | -10.469 | 8539 | 0.000 |
| Body Mass Index<br>(All: s, a, u ♂: a, e ♀: a) | 26.061 | 24.519 | 15.037 | 10771 | 0.000 |
| Intensity of liking dogs<br>(All: s, a, e, u ♂: e, u ♀: a, e, u) | 67.858 | 76.669 | -14.516 | 9482 | 0.000 |
| Intensity of liking cats<br>(All: s, a, e ♂: a, e ♀: e) | 61.225 | 71.854 | -15.926 | 9453 | 0.000 |
| Preference dogs to cats<br>(All: s, e, u ♂: u ♀: u) | 6.646 | 4.928 | 2.053 | 9391 | 0.040 |
| Sexual activity<br>(All: s, a, u ♂: a, u ♀: a, u) | -0.024 | 0.026 | -2.869 | 7373 | 0.004 |
| Sexual desire<br>(All: s, e ♂: a, e, u ♀: 0) | 7.266 | 6.975 | 6.806 | 7242 | 0.000 |

For the explanation of codes in the column 1 see the table 2.

In the sample, 2,044 (18.8%) subjects, namely 420 (9.83 %) men and 1,624 (24.7 %) women, were aware of their toxoplasmosis status. Among them, about 40.3% men and 34.3% women were tested in our lab during participation in various research projects, 49.4% men and 20.7% women were tested in relation to their health problems, 37.6% women were screened for toxoplasmosis in relation to their pregnancy and 10.4% men and 7.4% women were tested for other reasons. The seroprevalence of toxoplasmosis was much lower in men (18.3%, n =77) than in women (27.1%, n =440) (Chi^2^ = 13.6, p = 0.0002), which probably reflected opposite behavioral reaction of men and women to the infection (increased vs. decreased cooperativeness) [39] rather than real differences in the prevalence of toxoplasmosis in men and women [40].

### b) Association of keeping pets with wellbeing and health

Nearly all focal variables had highly irregular distributions and many of them correlated with sex, age, urbanization, and education (see the first column of the Tables 2-3). Therefore, we used nonparametric partial Kendall regression tests controlled for age, size of place of living, and education to search for associations between the animal-related variables and wellbeing and health, separately for male (Table 4) and female (Table 5) responders. For details, i.e. the correlations with particular source variables, see the Supplementary tables S1 and S2. The results showed positive effects of liking dogs on reported wellbeing (non-significant except the health domain in women, see S1-S2 Tables) and no effects of liking cats on wellbeing. Liking dogs had a positive effect on family situation in men and a negative effect on economic situation in women, while liking cats had a negative effect on economic situation in women. Liking dogs, and especially preferring dogs to cats, had positive effects on the mental health of both men and women. The effects on physical health were smaller, and in women were even negative. Liking cats had numerous and relatively strong negative effects on the mental health of both men and women and no effects on physical health. In both men and women, liking dogs had strong positive effects on sexual activity and sexual desire but very strong negative effects on the proxies of direct biological fitness and inclusive fitness – the number of children and siblings, respectively. The same was true for liking cats; here, however, the positive effect on sexual activity and sexual desire was not significant in men. Actually, the negative association of number of children with liking pets was stronger than with any other factors, including smoking, consuming alcohol, and (in women but not men) consuming illegal drugs.

**Table 4.**
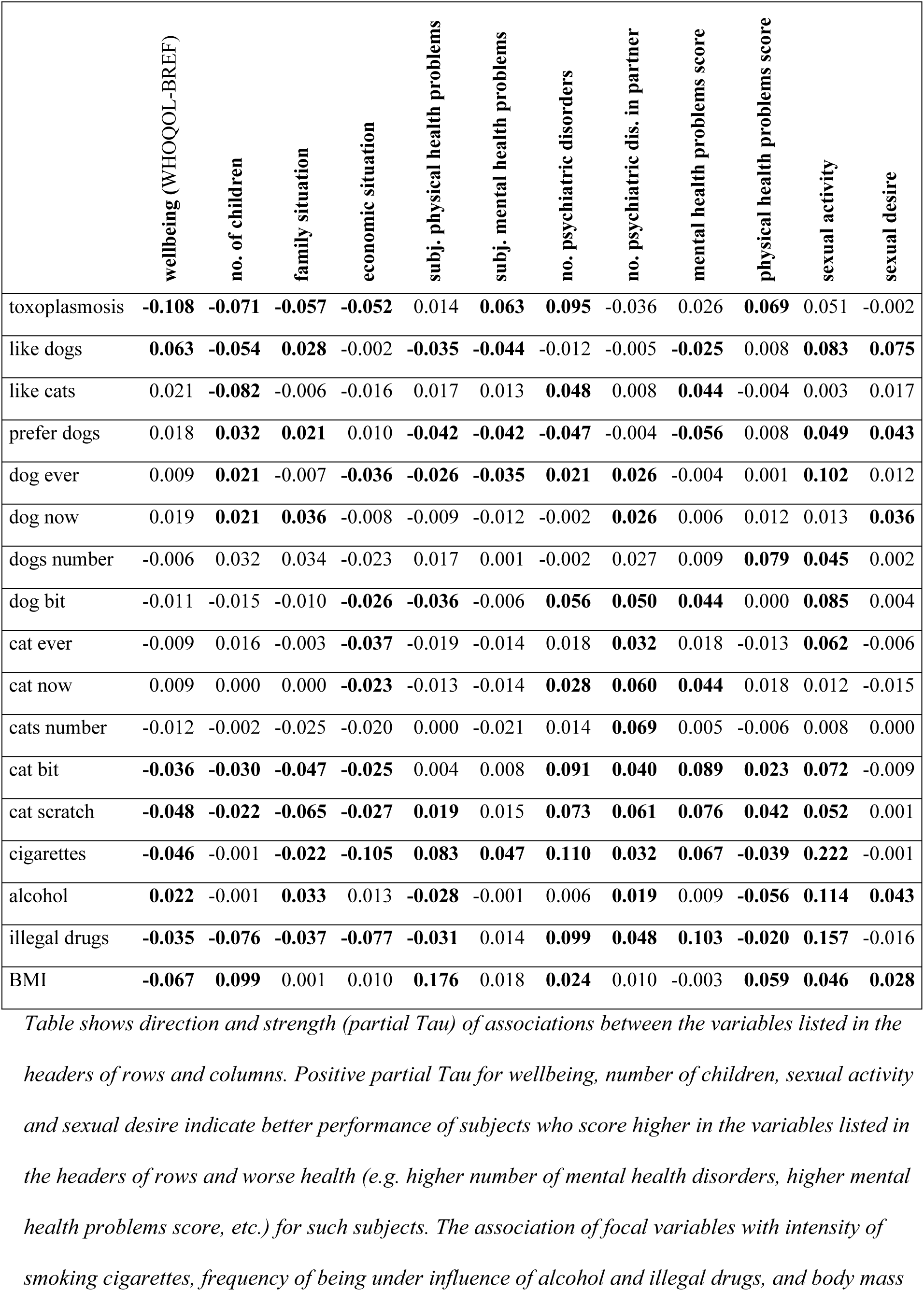
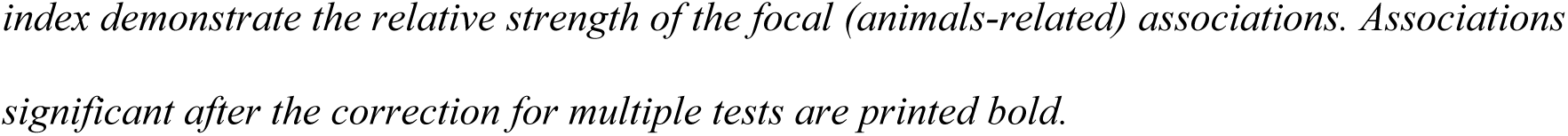
Association of animals-related variables with wellbeing and health of male responders

|  | wellbeing (WHOQOL-BREF) | no. of children | family situation | economic situation | subj. physical health problems | subj. mental health problems | no. psychiatric disorders | no. psychiatric dis. in partner | mental health problems score | physical health problems score | sexual activity | sexual desire |
| --- | --- | --- | --- | --- | --- | --- | --- | --- | --- | --- | --- | --- |
| toxoplasmosis | **-0.108** | **-0.071** | **-0.057** | **-0.052** | 0.014 | **0.063** | **0.095** | -0.036 | 0.026 | **0.069** | 0.051 | -0.002 |
| like dogs | **0.063** | **-0.054** | **0.028** | -0.002 | **-0.035** | **-0.044** | -0.012 | -0.005 | **-0.025** | 0.008 | **0.083** | **0.075** |
| like cats | 0.021 | **-0.082** | -0.006 | -0.016 | 0.017 | 0.013 | **0.048** | 0.008 | **0.044** | -0.004 | 0.003 | 0.017 |
| prefer dogs | 0.018 | **0.032** | **0.021** | 0.010 | **-0.042** | **-0.042** | **-0.047** | -0.004 | **-0.056** | 0.008 | **0.049** | **0.043** |
| dog ever | 0.009 | **0.021** | -0.007 | **-0.036** | **-0.026** | **-0.035** | **0.021** | **0.026** | -0.004 | 0.001 | **0.102** | 0.012 |
| dog now | 0.019 | **0.021** | **0.036** | -0.008 | -0.009 | -0.012 | -0.002 | **0.026** | 0.006 | 0.012 | 0.013 | **0.036** |
| dogs number | -0.006 | 0.032 | 0.034 | -0.023 | 0.017 | 0.001 | -0.002 | 0.027 | 0.009 | **0.079** | **0.045** | 0.002 |
| dog bit | -0.011 | -0.015 | -0.010 | **-0.026** | **-0.036** | -0.006 | **0.056** | **0.050** | **0.044** | 0.000 | **0.085** | 0.004 |
| cat ever | -0.009 | 0.016 | -0.003 | **-0.037** | -0.019 | -0.014 | 0.018 | **0.032** | 0.018 | -0.013 | **0.062** | -0.006 |
| cat now | 0.009 | 0.000 | 0.000 | **-0.023** | -0.013 | -0.014 | **0.028** | **0.060** | **0.044** | 0.018 | 0.012 | -0.015 |
| cats number | -0.012 | -0.002 | -0.025 | -0.020 | 0.000 | -0.021 | 0.014 | **0.069** | 0.005 | -0.006 | 0.008 | 0.000 |
| cat bit | **-0.036** | **-0.030** | **-0.047** | **-0.025** | 0.004 | 0.008 | **0.091** | **0.040** | **0.089** | **0.023** | **0.072** | -0.009 |
| cat scratch | **-0.048** | **-0.022** | **-0.065** | **-0.027** | **0.019** | 0.015 | **0.073** | **0.061** | **0.076** | **0.042** | **0.052** | 0.001 |
| cigarettes | **-0.046** | -0.001 | **-0.022** | **-0.105** | **0.083** | **0.047** | **0.110** | **0.032** | **0.067** | **-0.039** | **0.222** | -0.001 |
| alcohol | **0.022** | -0.001 | **0.033** | 0.013 | **-0.028** | -0.001 | 0.006 | **0.019** | 0.009 | **-0.056** | **0.114** | **0.043** |
| illegal drugs | **-0.035** | **-0.076** | **-0.037** | **-0.077** | **-0.031** | 0.014 | **0.099** | **0.048** | **0.103** | **-0.020** | **0.157** | -0.016 |
| BMI | **-0.067** | **0.099** | 0.001 | 0.010 | **0.176** | 0.018 | **0.024** | 0.010 | -0.003 | **0.059** | **0.046** | **0.028** |

*Table shows direction and strength (partial Tau) of associations between the variables listed in the* *headers of rows and columns. Positive partial Tau for wellbeing, number of children, sexual activity* *and sexual desire indicate better performance of subjects who score higher in the variables listed in* *the headers of rows and worse health (e.g. higher number of mental health disorders, higher mental* *health problems score, etc.) for such subjects. The association of focal variables with intensity of* *smoking cigarettes, frequency of being under influence of alcohol and illegal drugs, and body mass*
*index demonstrate the relative strength of the focal (animals-related) associations. Associations significant after the correction for multiple tests are printed bold.*

**Table 5.** Association of animal-related variables with wellbeing and health of female responders

|  | wellbeing (WHOQOL-BREF) | no. of children | family situation | economic situation | subj. physical health problems | subj. mental health problems | no. psychiatric disorders | no. psychiatric dis. in partner | mental health problems score | physical health problems score | sexual activity | sexual desire |
| --- | --- | --- | --- | --- | --- | --- | --- | --- | --- | --- | --- | --- |
| toxoplasmosis | -0.016 | -0.006 | **-0.035** | **-0.049** | -0.013 | **0.042** | **0.053** | -0.017 | **0.056** | **0.090** | -0.030 | **-0.066** |
| like dogs | **0.019** | **-0.095** | **0.017** | **-0.023** | -0.013 | -0.005 | 0.000 | 0.000 | **-0.019** | **0.021** | **0.038** | **0.103** |
| like cats | 0.007 | **-0.080** | -0.001 | **-0.032** | 0.010 | 0.004 | **0.055** | **0.019** | **0.044** | 0.013 | **0.041** | **0.033** |
| prefer dogs | 0.009 | -0.001 | 0.008 | 0.013 | **-0.025** | -0.008 | **-0.046** | -0.013 | **-0.043** | 0.004 | -0.003 | **0.045** |
| dog ever | **-0.019** | **0.020** | **-0.036** | **-0.045** | -0.012 | -0.002 | **0.030** | **0.025** | **0.022** | 0.008 | **0.072** | -0.013 |
| dog now | -0.006 | **-0.026** | -0.009 | **-0.029** | -0.002 | -0.016 | 0.013 | 0.012 | 0.005 | -0.004 | 0.016 | 0.013 |
| dogs number | **-0.046** | **-0.027** | -0.013 | **-0.034** | 0.019 | 0.012 | 0.014 | -0.010 | 0.019 | 0.004 | **0.038** | **0.058** |
| dog bit | **-0.042** | -0.009 | **-0.033** | **-0.046** | **-0.019** | 0.004 | **0.070** | 0.003 | **0.072** | **0.030** | **0.052** | **0.015** |
| cat ever | **-0.034** | **0.029** | **-0.018** | **-0.060** | 0.008 | -0.001 | **0.055** | **0.041** | **0.040** | 0.010 | **0.047** | **-0.026** |
| cat now | **-0.016** | **-0.023** | **-0.014** | **-0.041** | **0.018** | -0.010 | **0.060** | **0.013** | **0.053** | 0.009 | **0.029** | **-0.025** |
| cats number | **-0.044** | **-0.040** | -0.009 | **-0.057** | **0.029** | 0.016 | **0.030** | **0.031** | **0.033** | 0.018 | **0.026** | **-0.031** |
| cat bit | **-0.044** | **-0.056** | **-0.028** | **-0.043** | 0.007 | **0.018** | **0.085** | **0.014** | **0.083** | **0.022** | **0.056** | **-0.024** |
| cat scratch | **-0.063** | **-0.057** | **-0.034** | **-0.050** | **0.038** | **0.034** | **0.074** | -0.004 | **0.100** | **0.040** | **0.042** | **-0.041** |
| cigarettes | **-0.034** | **-0.046** | **-0.066** | **-0.079** | **0.034** | **0.030** | **0.105** | **0.029** | **0.084** | -0.011 | **0.248** | **0.028** |
| alcohol | **0.014** | **-0.077** | **-0.016** | **-0.018** | **-0.045** | 0.009 | **0.024** | 0.010 | **0.052** | **-0.023** | **0.150** | **0.023** |
| illegal drugs | **-0.032** | **-0.058** | **-0.035** | **-0.049** | -0.004 | 0.014 | **0.067** | **0.044** | **0.075** | 0.010 | **0.158** | 0.006 |
| BMI | **-0.064** | **0.066** | **-0.031** | **-0.042** | **0.222** | **0.026** | **0.051** | **0.034** | **0.032** | **0.072** | **0.021** | -0.001 |

*For legend, see the table 4.*

In men, keeping dogs, currently keeping and especially having ever kept, had positive effects on some domains of wellbeing (social relationships, and psychological domains); however, it had a negative effect on economic situation. Number of dogs correlated negatively with health and environment domains, with total wellbeing score, and economic situation. In women, keeping dogs had a negative effect on the environment domain of wellbeing and economic situation, and the number of dogs in a house correlated negatively with health and environment domains, total wellbeing score, and economic situation. Keeping cats and number of cats in a house had minimum effects on the wellbeing of men, except a negative effect on economic situation; however, it had numerous negative effects (especially the number of cats in house and ever keeping a cat) on wellbeing and economic situation of women. The effects of keeping pets on wellbeing depended, both in strength and direction, on the age of subjects; stronger positive effects were observed only for older participants (Figs 1-2, S1-S4 Tables).

**Fig. 1.**
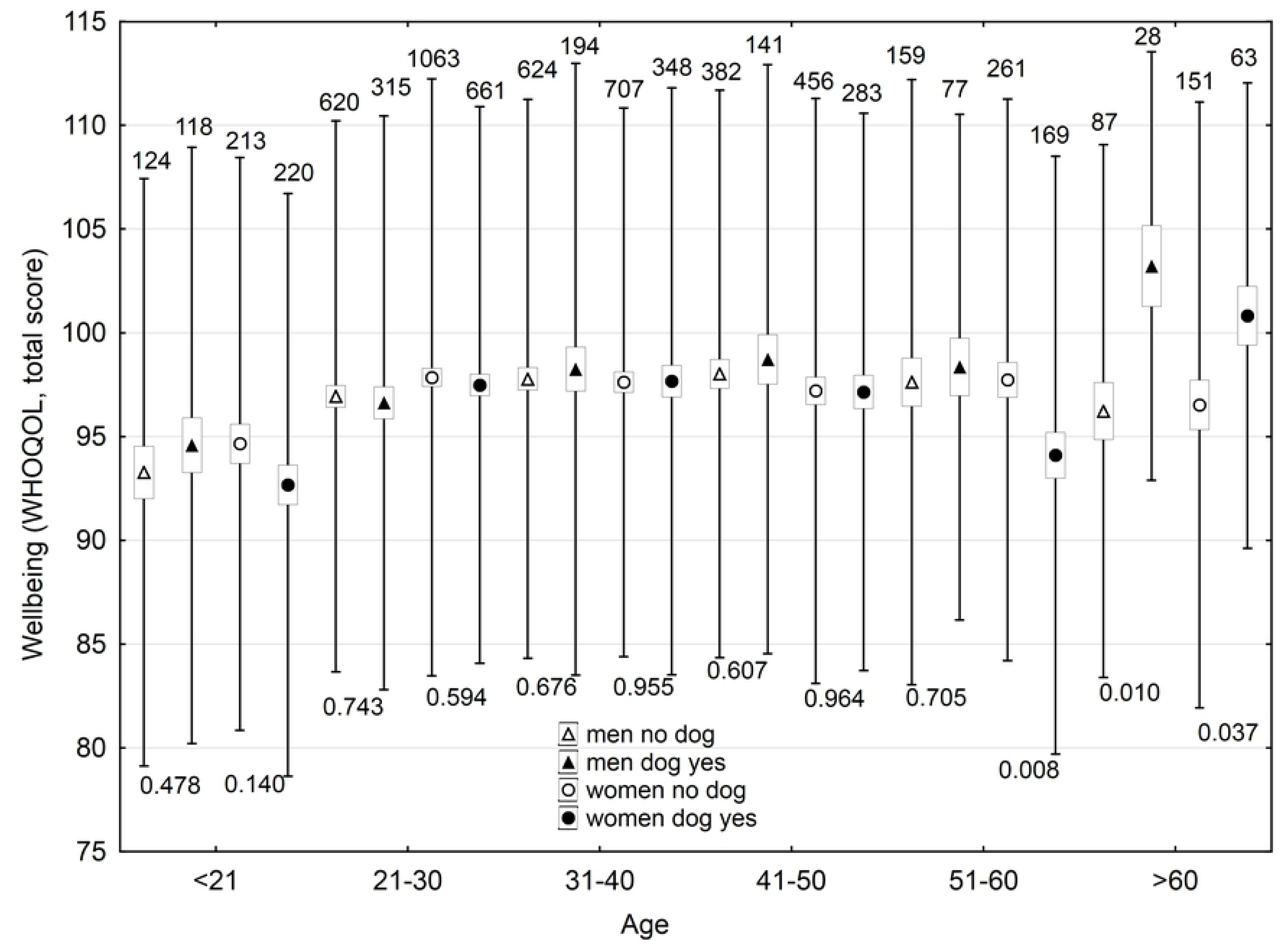
Effect of keeping a dog now on wellbeing of men and women of different age. The boxes, spreads, upper numbers and lower numbers show standard errors, standard deviations, numbers of subjects in particular category and p-values of two-sided t-tests, respectively.

**Fig. 2.**
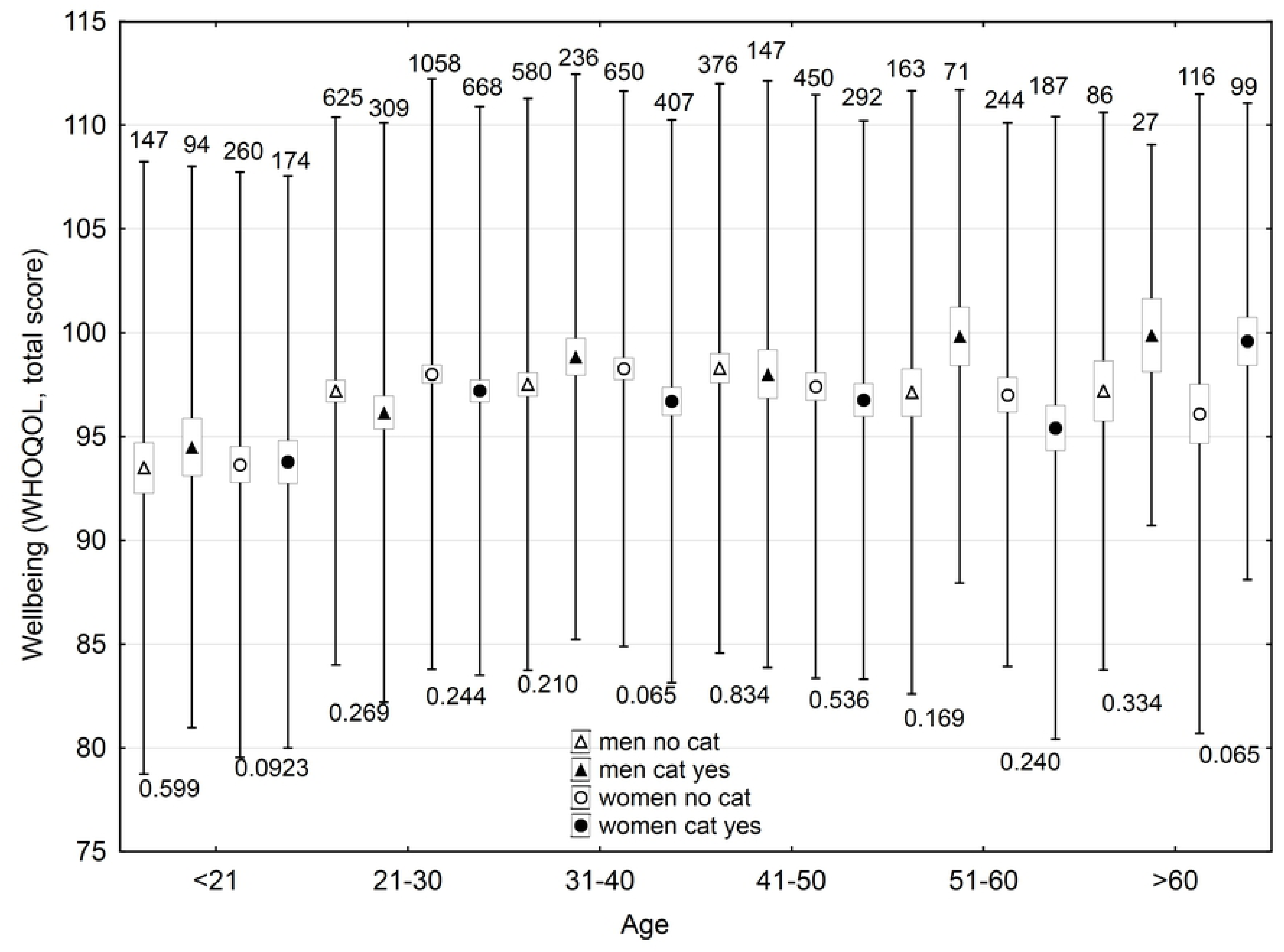
Effect of keeping a cat now on wellbeing of men and women of different age. The boxes, spreads, upper numbers and lower numbers show standard errors, standard deviations, numbers of subjects in particular category and p-values of two-sided t-tests, respectively.

In men, keeping a dog had small, mostly non-significant effects on mental and physical health; however, number of dogs in a house had a negative effect on physical health. In women, ever having kept a dog, but not keeping a dog now, or number of dogs in a house had negative effects on mental health. Ever keeping a dog also correlated positively with sexual activity, and the number of dogs correlated with both sexual activity and sexual desire. In men, keeping a cat now correlated negatively with mental health and the number of cats in a house correlated positively with the number of undiagnosed mental health disorders in their partners. Ever having kept a cat correlated positively with sexual activity; however, no correlation was observed with sexual desire or number of children. In women, keeping a cat and number of cats correlated positively with mental health problems (especially with the number of diagnosed and undiagnosed mental health disorders). It correlated positively with sexual activity and negatively with sexual desire. Ever having kept a cat correlated positively, while keeping a cat now and number of cats in a house negatively, with number of children.

### c) Association of animal-related injuries with wellbeing and health

The most numerous (strong and negative) effects were observed between focal variables and animal-(especially cat-) related injuries (Figs. 3-8 and supplementary Figs. S5-S13). In men, dog-related injuries had negative effects on the environmental domain of wellbeing, economic situation, and mental health, and a positive effect on sexual activity, while in women it had significant negative effects on all four domains of wellbeing, total wellbeing score, family situation, economic situation, and mental and physical health. It also had a positive effect on sexual activity. In men, cat-related injury, especially cat scratching, had strong negative effects on all domains of wellbeing and total wellbeing score, family situation, economic situation, mental and physical health and number of children. It had a positive effect on sexual activity and no effect on sexual desire. In women, we observed the same effects as in men, only stronger. Women with animal-related injuries also had lower numbers of siblings and lower sexual desire. The effects of animal-related injuries were of similar strength as those of four risk factors used as the internal control. They were stronger and more negative than that of consuming alcohol, weaker than that of consuming illegal drugs and approximately the same as smoking and high BMI.

**Fig. 3.**
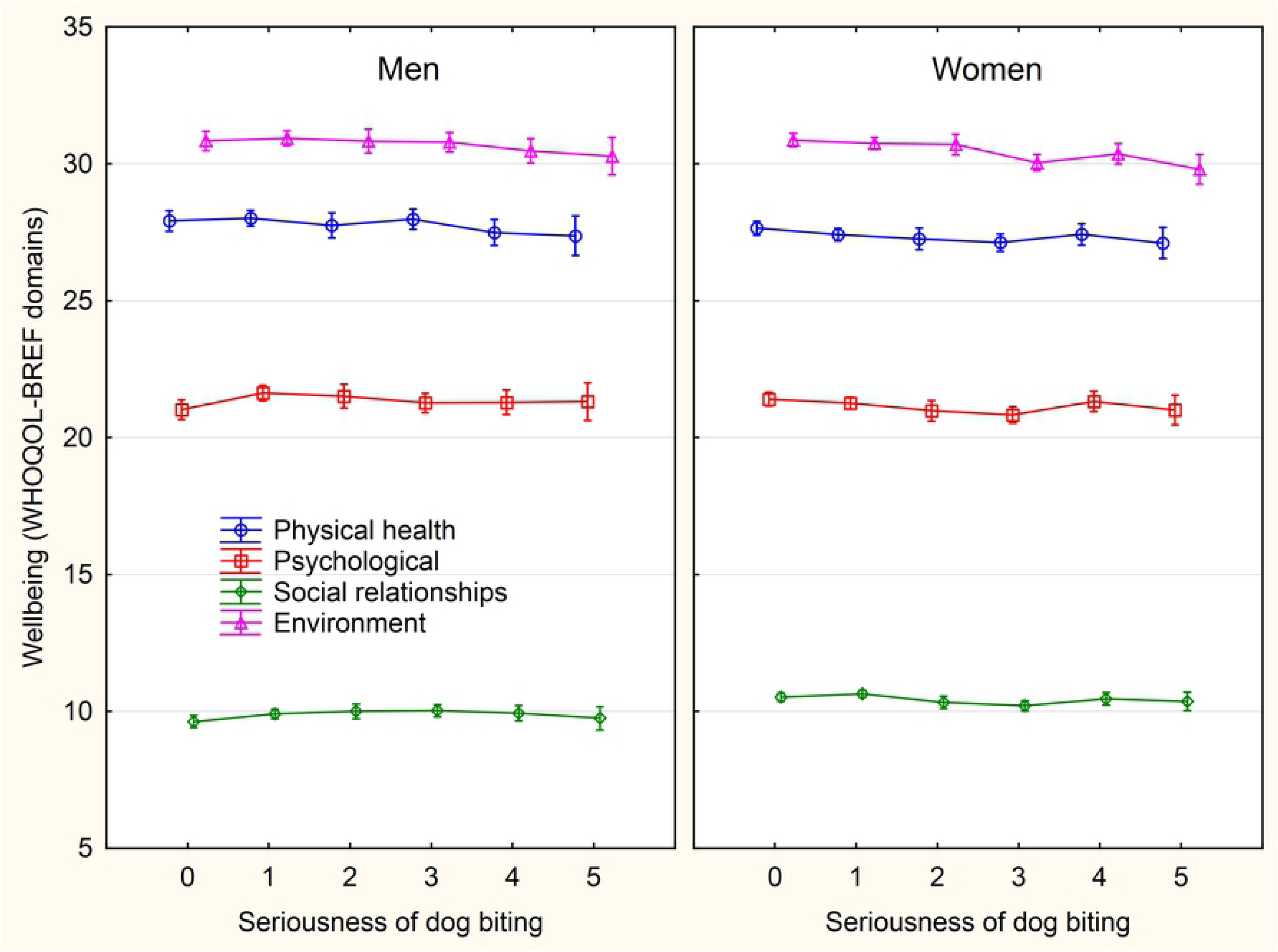
Association between intensity of sustained dog biting and wellbeing – the scores of WHOQOL-BREF domains. The categories on x-axis describing the intensity of being injured by a pet are: 0-never, 1-only while playing, 2-only as a warning, 3-yes, minor injury (only skin cut), 4-yes, moderate injury (bleeding), 5-yes, serious injury, I had to seek medical treatment.

**Fig. 4.**
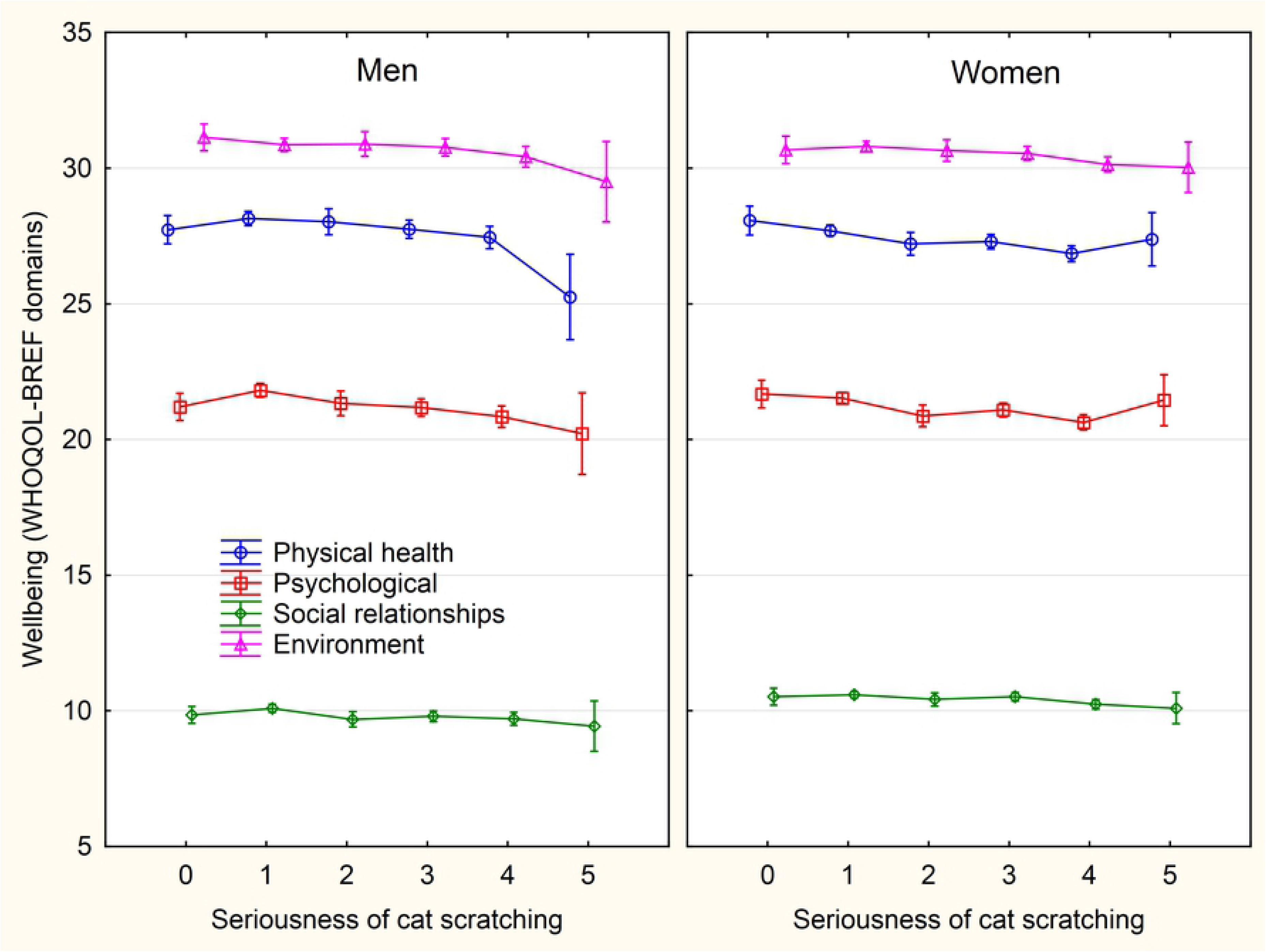
Association between intensity of sustained cat scratching and wellbeing – the scores of WHOQOL-BREF domains. The categories on x-axis describing the intensity of being injured by a pet are: 0-never, 1-only while playing, 2-only as a warning, 3-yes, minor injury (only skin cut), 4-yes, moderate injury (bleeding), 5-yes, serious injury, I had to seek medical treatment.

**Fig. 5.**
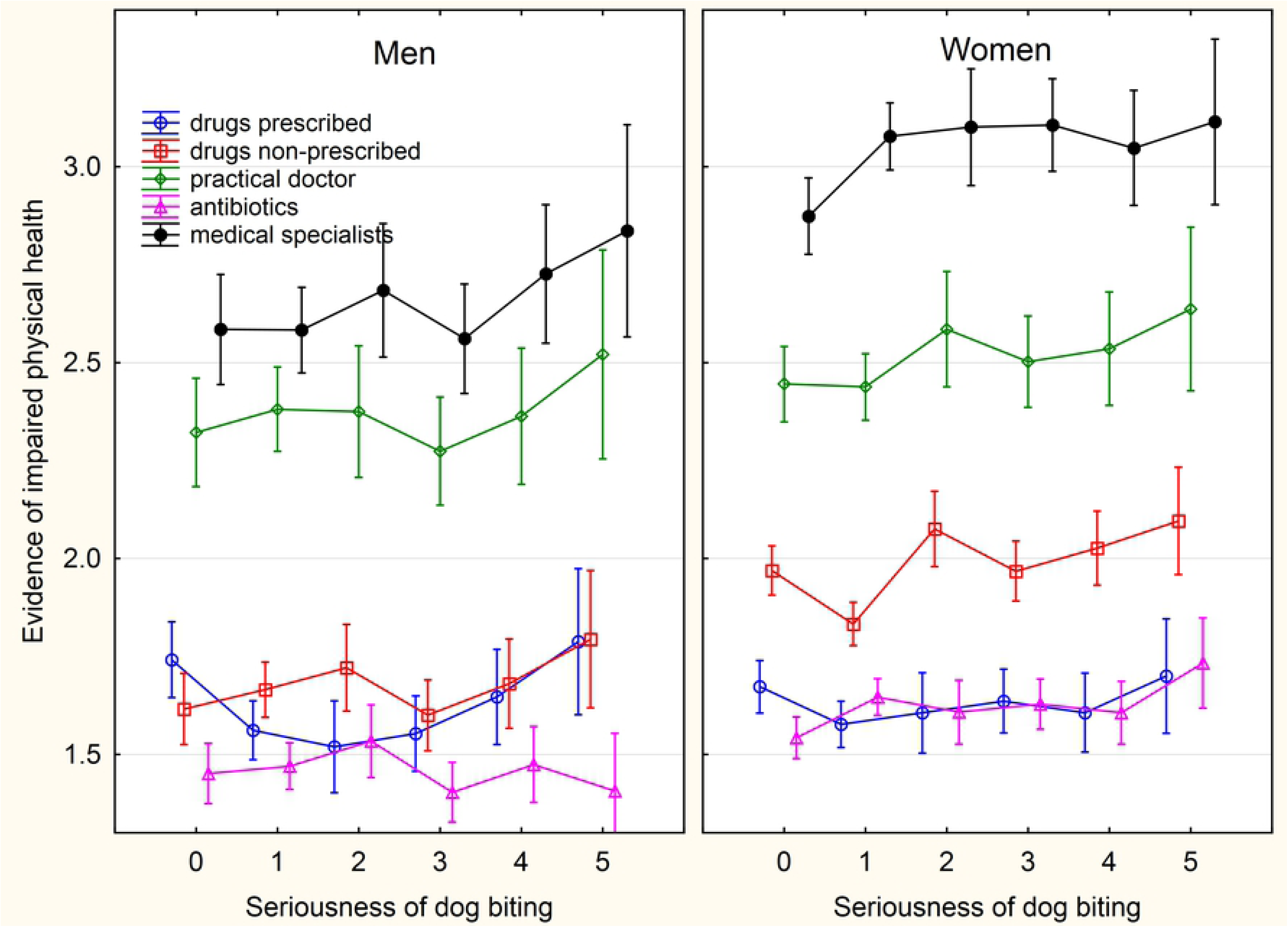
Association between intensity of sustained dog biting and physical health related variables. The categories on x-axis describing the intensity of being injured by a pet are: 0-never, 1-only while playing, 2-only as a warning, 3-yes, minor injury (only skin cut), 4-yes, moderate injury (bleeding), 5-yes, serious injury, I had to seek medical treatment.

**Fig. 6.**
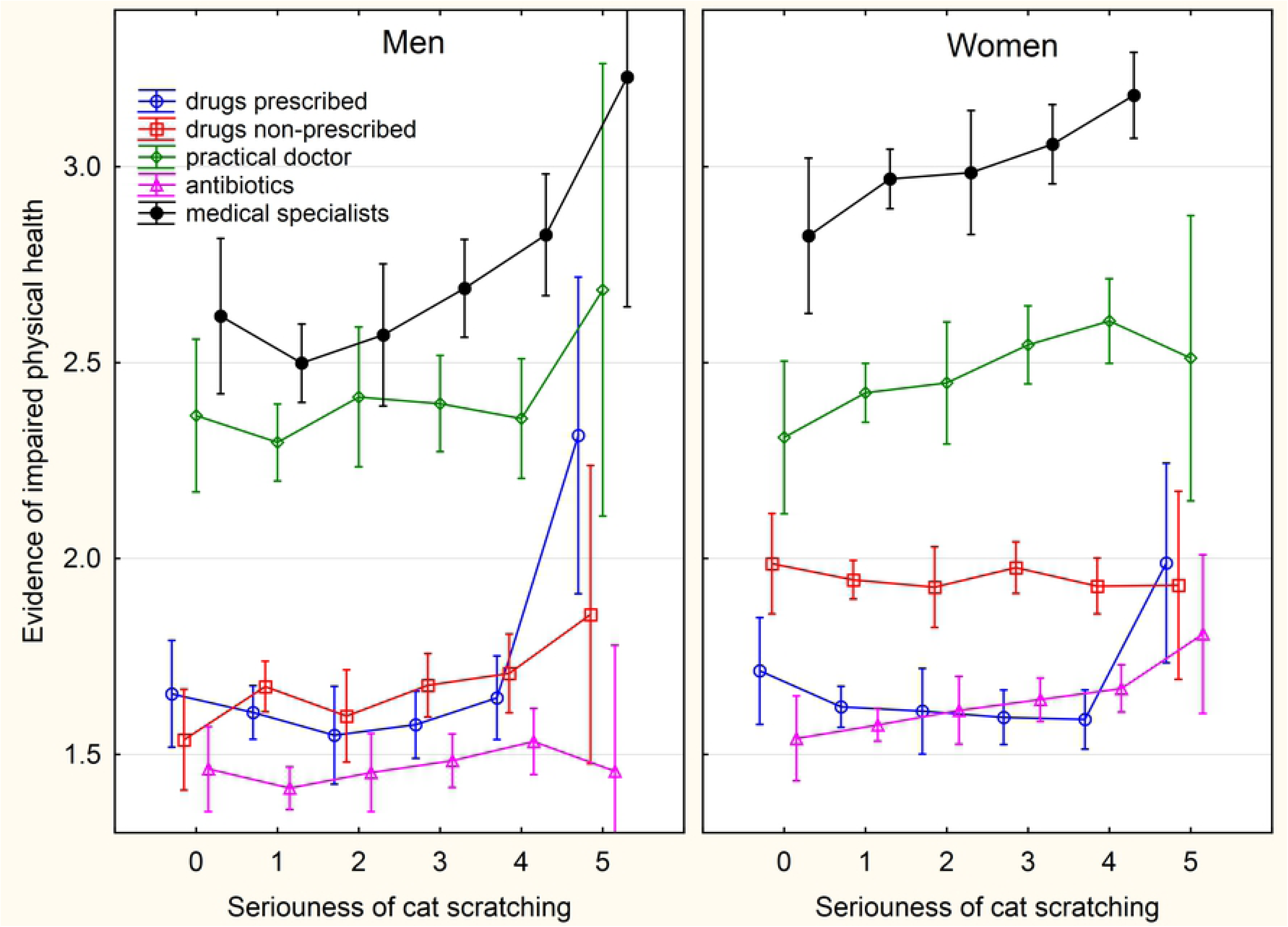
Association between intensity of sustained cat scratching and physical health related variables. The categories on x-axis describing the intensity of being injured by a pet are: 0-never, 1-only while playing, 2-only as a warning, 3-yes, minor injury (only skin cut), 4-yes, moderate injury (bleeding), 5-yes, serious injury, I had to seek medical treatment.

**Fig. 7.**
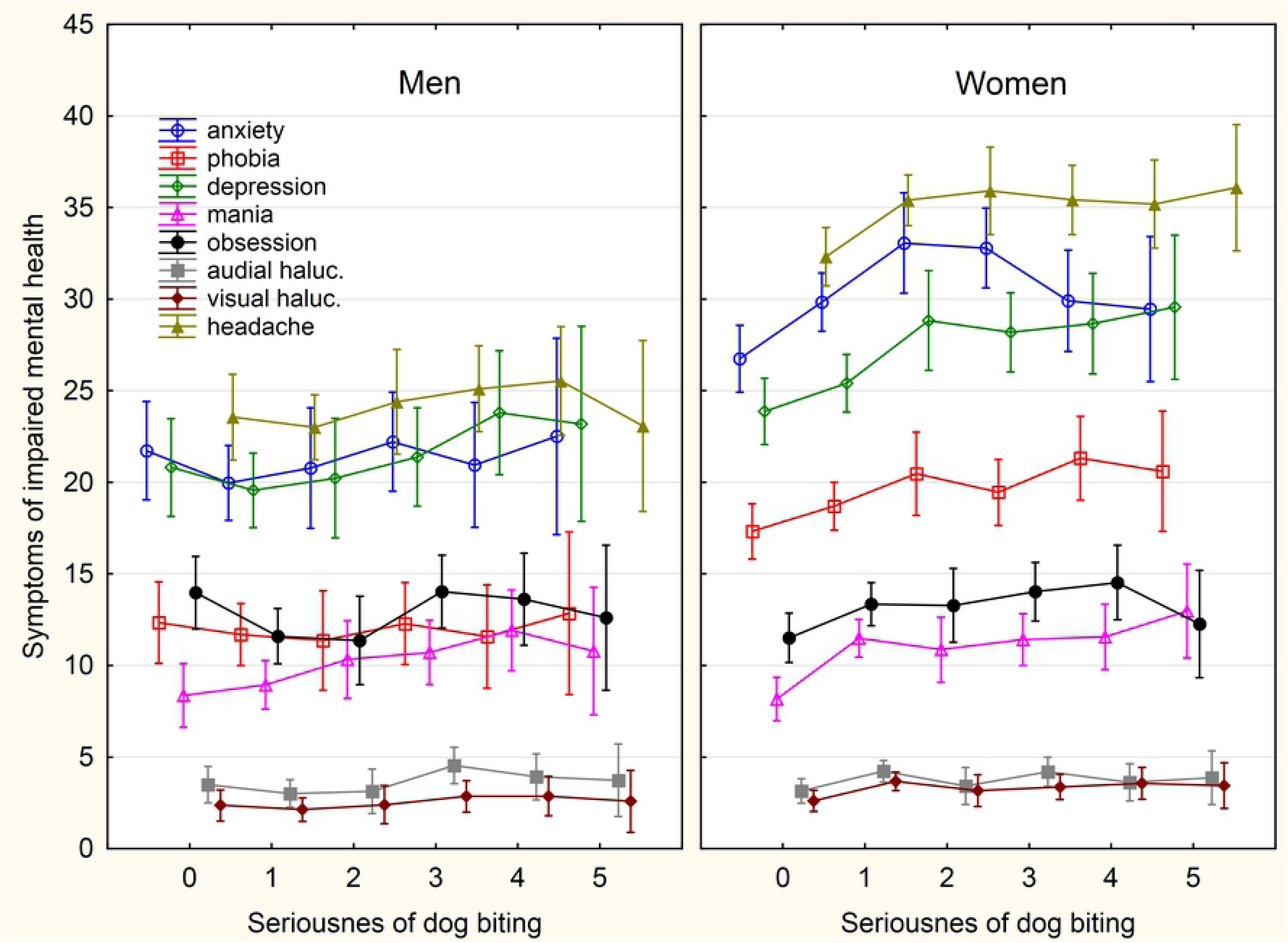
Association between intensity of sustained dog biting and symptoms of impaired mental health. The categories on x-axis describing the intensity of being injured by a pet are: 0-never, 1-only while playing, 2-only as a warning, 3-yes, minor injury (only skin cut), 4-yes, moderate injury (bleeding), 5-yes, serious injury, I had to seek medical treatment.

**Fig. 8.**
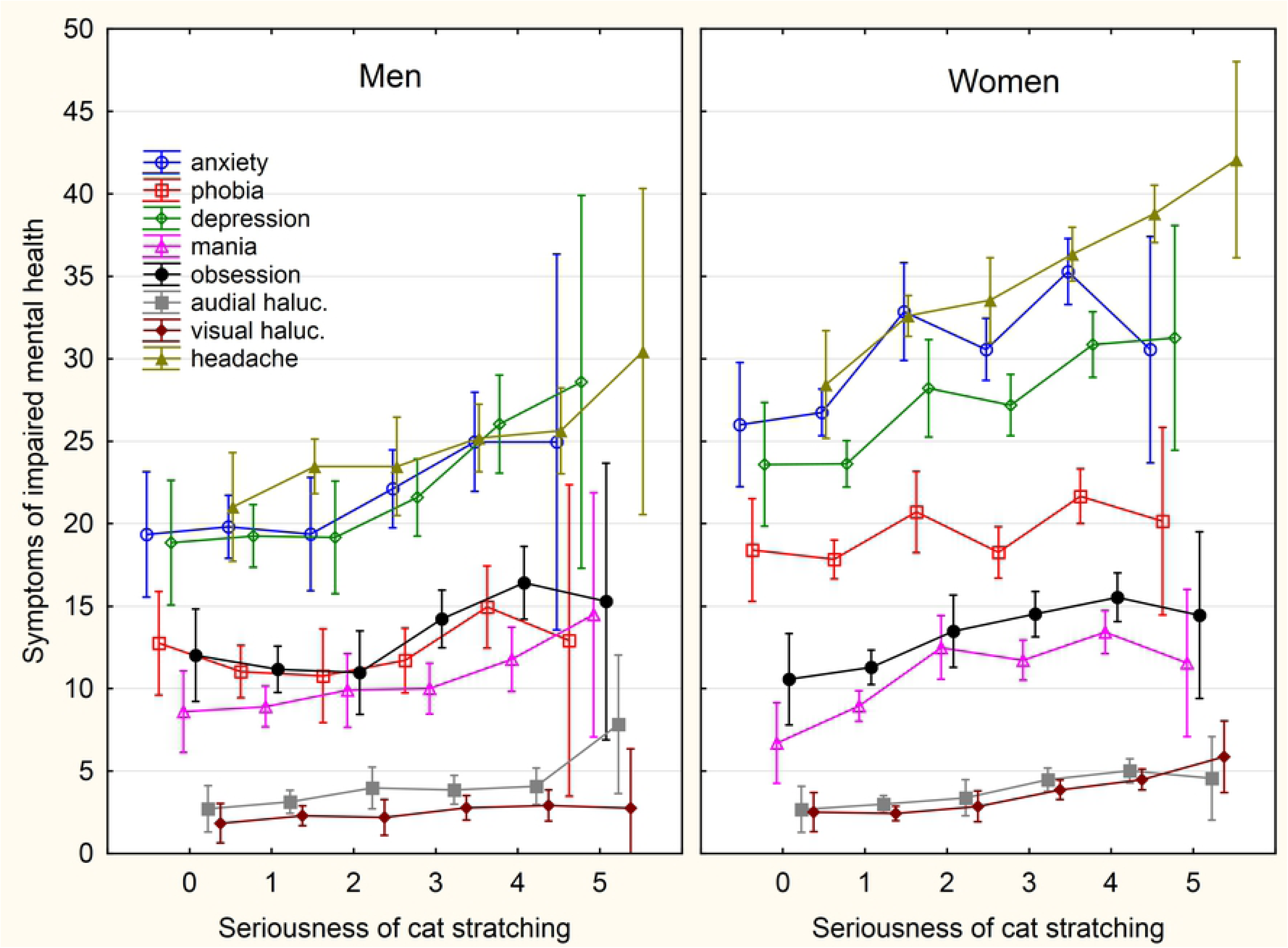
Association between intensity of sustained cat scratching and symptoms of impaired mental health. The categories on x-axis describing the intensity of being injured by a pet are: 0-never, 1-only while playing, 2-only as a warning, 3-yes, minor injury (only skin cut), 4-yes, moderate injury (bleeding), 5-yes, serious injury, I had to seek medical treatment

Latent infection with the parasite *Toxoplasma* had strong effects on wellbeing and physical health, even in comparison with cat-related injuries and consumption of illegal drugs. Due to the relatively low number of infected men (77), not all correlations, however, were statistically significant in this group. In men, toxoplasmosis had strong negative effects on all domains of wellbeing except social relationships, on the total wellbeing score, family situation and economic situation. It also had negative effects on physical and mental health, and number of children. A positive relation was observed with sexual activity and especially with number of siblings. In women, a negative correlation existed between toxoplasmosis and the environment domain of wellbeing, family situation and economic situation, with physical health, mental health, and sexual activity and sexual desire.

### d) Is keeping pets responsible for the association of liking pets with wellbeing and health, and are animal-related injuries or toxoplasmosis responsible for the association of keeping pets with wellbeing and health?

People who like cats and dogs also have a higher probability of keeping them. To check whether the positive association of liking dogs and wellbeing and negative association between liking cats and dogs and mental health also existed in subjects who never kept these animals, we repeated all analyses on the subsets of men and women who never kept a dog or a cat. The results shown in the Supplementary tables S3-S6 were similar to those obtained by the analysis of the whole population, suggesting that keeping dogs and cats is not responsible for most of the observed associations between liking animals and wellbeing- and health-related variables. Similarly, the negative associations between keeping animals and mental health could theoretically just be a secondary effect of much stronger negative association between sustaining animal-related injuries and mental and physical health problems. Again, the pattern of association was very similar for the subjects who reported not to have been injured by cats and dogs (Supplementary tables S7-S10).

It is known, and also our current data strongly suggests, that latent infection with the parasite *Toxoplasma gondii* is associated with many mental health disorders and with impaired mental and physical health in general [41, 42]. Contact with cats (or rather the garden soil containing cat feces with *Toxoplasma* oocysts) and in some studies, also keeping dogs, is considered to be an important source of the *Toxoplasma* infection [40]. Multivariate logistic regression showed that *Toxoplasma* seropositivity was positively related to female sex (O.R. 1.68, C.I._95_ 1.24-2.26), age (O.R._range_ 3.28, C.I._95_ 1.24-2.26), intensity of being bitten by a cat (O.R._range_ 2.47, C.I._95_ 1.51-4.03), and intensity of being bitten by a dog (O.R._range_ 1.99, C.I._95_ 1.39-2.86). It was negatively but not significantly associated with urbanization (O.R._range_ 0.60, C.I._95_ 0.44-2.82). Also, the associations of *Toxoplasma* infection with education, keeping a dog and keeping a cat or with being scratched by a cat were nonsignificant (p > 0.80). The partial Kendal correlation analyses performed on the subsets of *Toxoplasma* seronegative subjects, namely 343 men and 1,184 women, showed that nearly all associations observed on these subsets were much stronger than the same associations observed in the whole set (Tables 6, 7, S11, S12). This suggests that toxoplasmosis was not responsible for the observed associations between the animal-related variables and human wellbeing, health, and fitness.

**Table 6.**
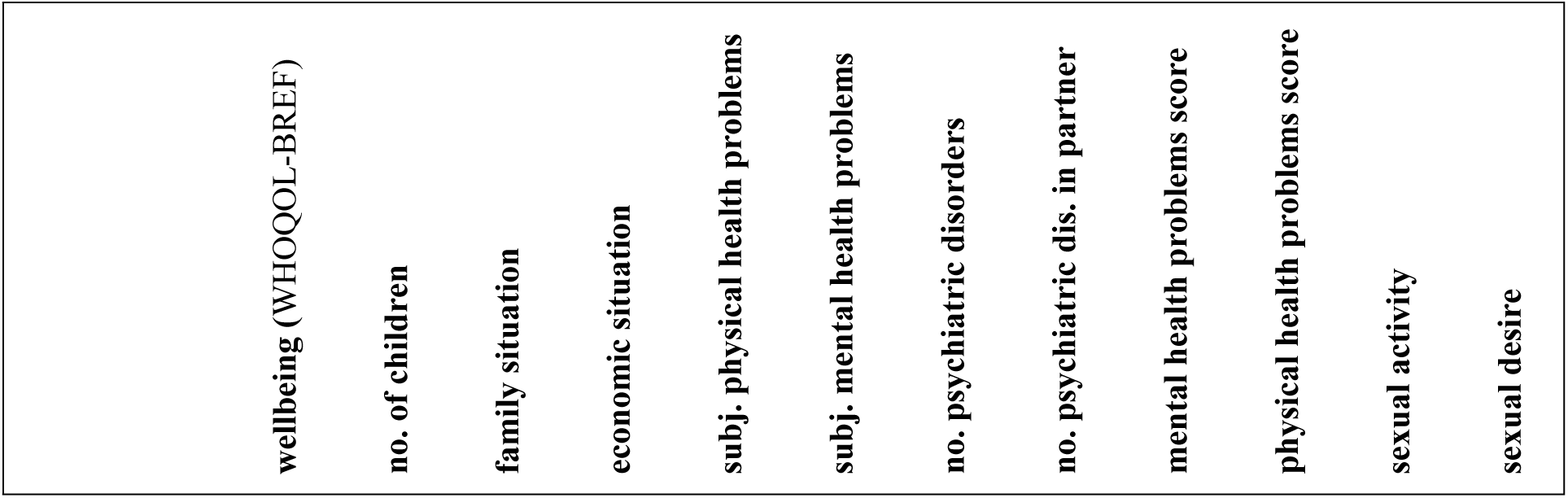
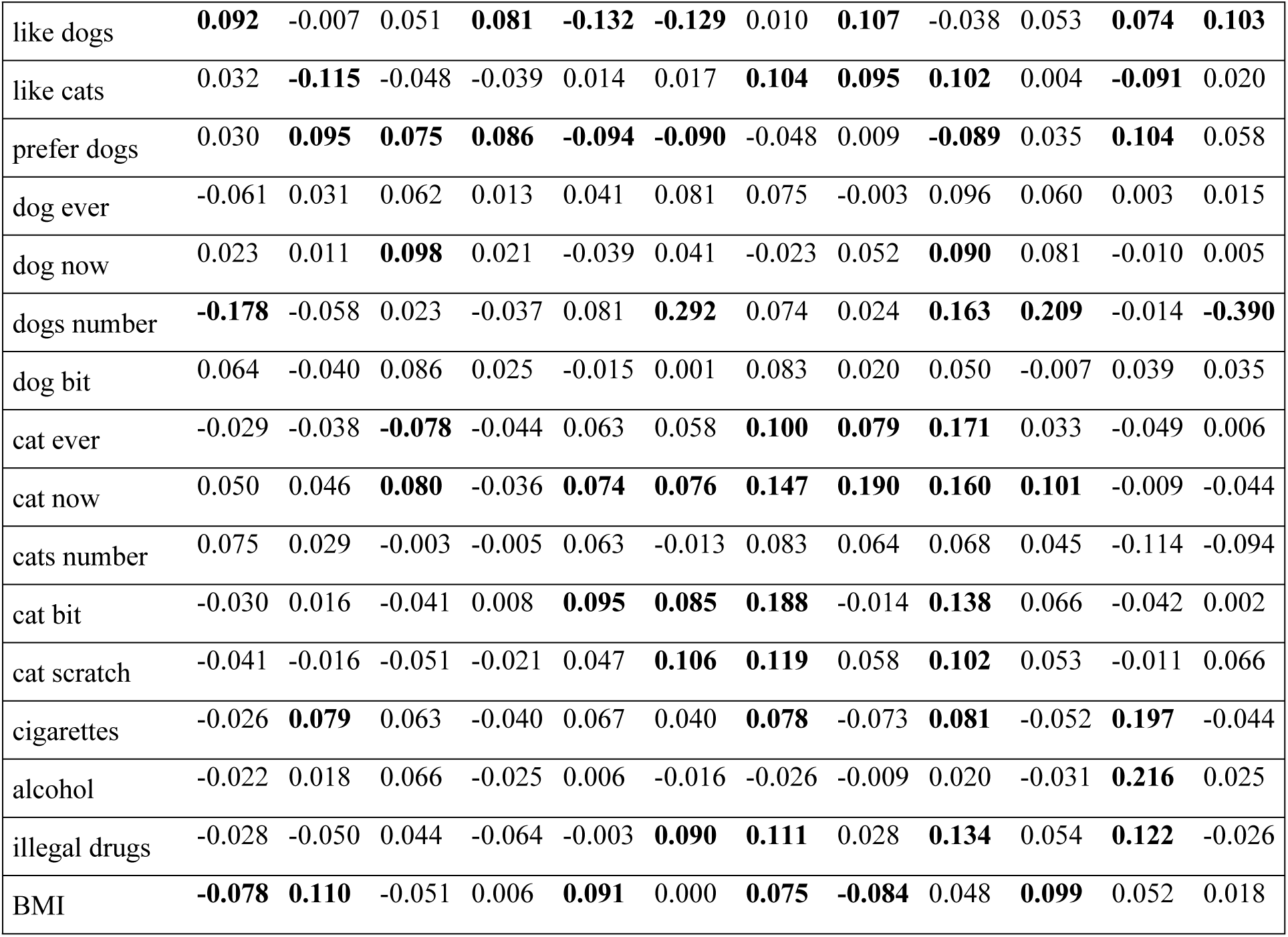
Association of animals-related variables with wellbeing and health of *Toxoplasma*-free male responders

| wellbeing (WHOQOL-BREF) | no. of children | family situation | economic situation | subj. physical health problems | subj. mental health problems | no. psychiatric disorders | no. psychiatric dis. in partner | mental health problems score | physical health problems score | sexual activity | sexual desire |
| --- | --- | --- | --- | --- | --- | --- | --- | --- | --- | --- | --- |

|  |  |  |  |  |  |  |  |  |  |  |  |  |
| --- | --- | --- | --- | --- | --- | --- | --- | --- | --- | --- | --- | --- |
| like dogs | **0.092** | -0.007 | 0.051 | **0.081** | **-0.132** | **-0.129** | 0.010 | **0.107** | -0.038 | 0.053 | **0.074** | **0.103** |
| like cats | 0.032 | **-0.115** | -0.048 | -0.039 | 0.014 | 0.017 | **0.104** | **0.095** | **0.102** | 0.004 | **-0.091** | 0.020 |
| prefer dogs | 0.030 | **0.095** | **0.075** | **0.086** | **-0.094** | **-0.090** | -0.048 | 0.009 | **-0.089** | 0.035 | **0.104** | 0.058 |
| dog ever | -0.061 | 0.031 | 0.062 | 0.013 | 0.041 | 0.081 | 0.075 | -0.003 | 0.096 | 0.060 | 0.003 | 0.015 |
| dog now | 0.023 | 0.011 | **0.098** | 0.021 | -0.039 | 0.041 | -0.023 | 0.052 | **0.090** | 0.081 | -0.010 | 0.005 |
| dogs number | **-0.178** | -0.058 | 0.023 | -0.037 | 0.081 | **0.292** | 0.074 | 0.024 | **0.163** | **0.209** | -0.014 | **-0.390** |
| dog bit | 0.064 | -0.040 | 0.086 | 0.025 | -0.015 | 0.001 | 0.083 | 0.020 | 0.050 | -0.007 | 0.039 | 0.035 |
| cat ever | -0.029 | -0.038 | **-0.078** | -0.044 | 0.063 | 0.058 | **0.100** | **0.079** | **0.171** | 0.033 | -0.049 | 0.006 |
| cat now | 0.050 | 0.046 | **0.080** | -0.036 | **0.074** | **0.076** | **0.147** | **0.190** | **0.160** | **0.101** | -0.009 | -0.044 |
| cats number | 0.075 | 0.029 | -0.003 | -0.005 | 0.063 | -0.013 | 0.083 | 0.064 | 0.068 | 0.045 | -0.114 | -0.094 |
| cat bit | -0.030 | 0.016 | -0.041 | 0.008 | **0.095** | **0.085** | **0.188** | -0.014 | **0.138** | 0.066 | -0.042 | 0.002 |
| cat scratch | -0.041 | -0.016 | -0.051 | -0.021 | 0.047 | **0.106** | **0.119** | 0.058 | **0.102** | 0.053 | -0.011 | 0.066 |
| cigarettes | -0.026 | **0.079** | 0.063 | -0.040 | 0.067 | 0.040 | **0.078** | -0.073 | **0.081** | -0.052 | **0.197** | -0.044 |
| alcohol | -0.022 | 0.018 | 0.066 | -0.025 | 0.006 | -0.016 | -0.026 | -0.009 | 0.020 | -0.031 | **0.216** | 0.025 |
| illegal drugs | -0.028 | -0.050 | 0.044 | -0.064 | -0.003 | **0.090** | **0.111** | 0.028 | **0.134** | 0.054 | **0.122** | -0.026 |
| BMI | **-0.078** | **0.110** | -0.051 | 0.006 | **0.091** | 0.000 | **0.075** | **-0.084** | 0.048 | **0.099** | 0.052 | 0.018 |

**Table 7.**
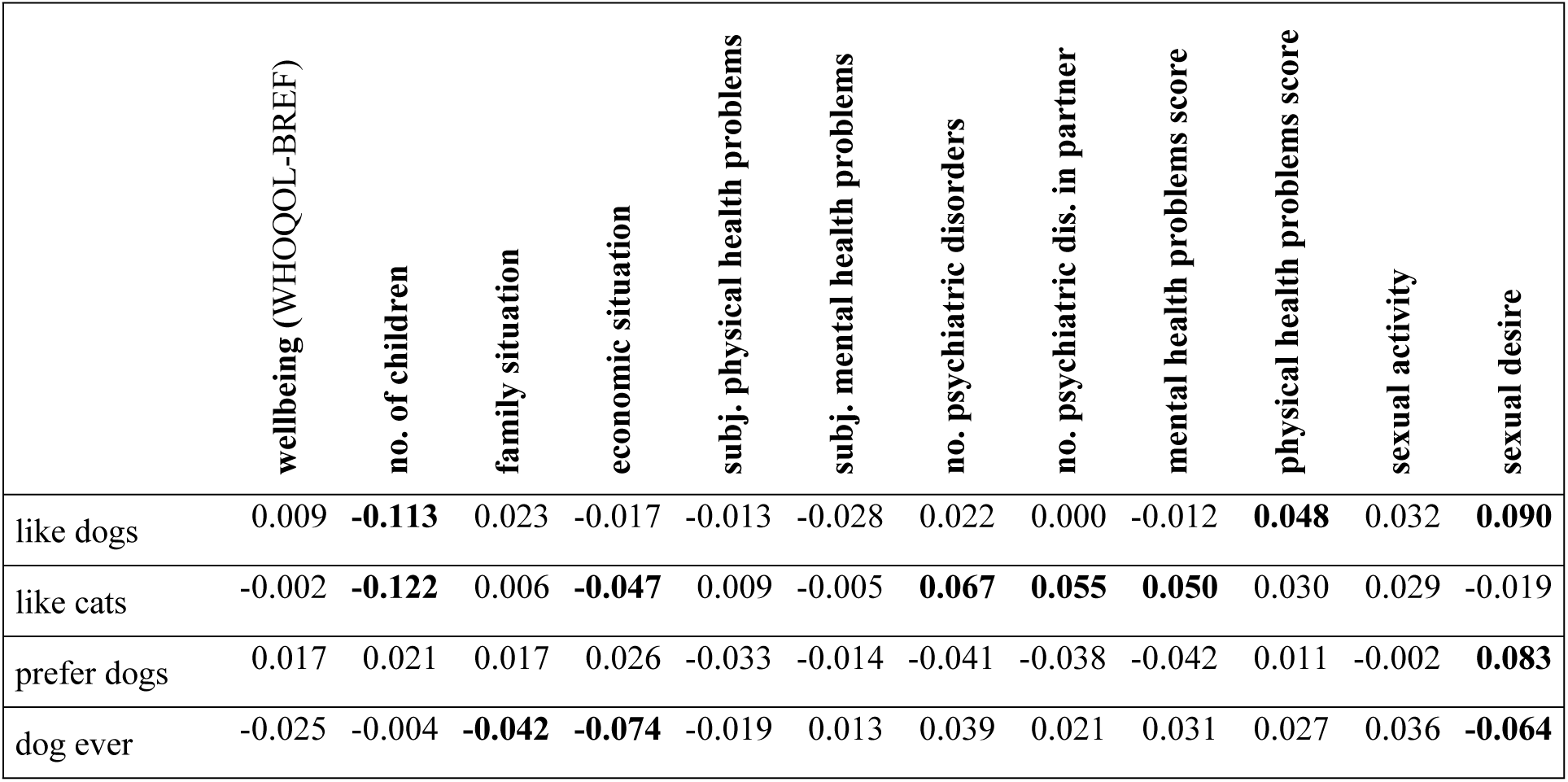
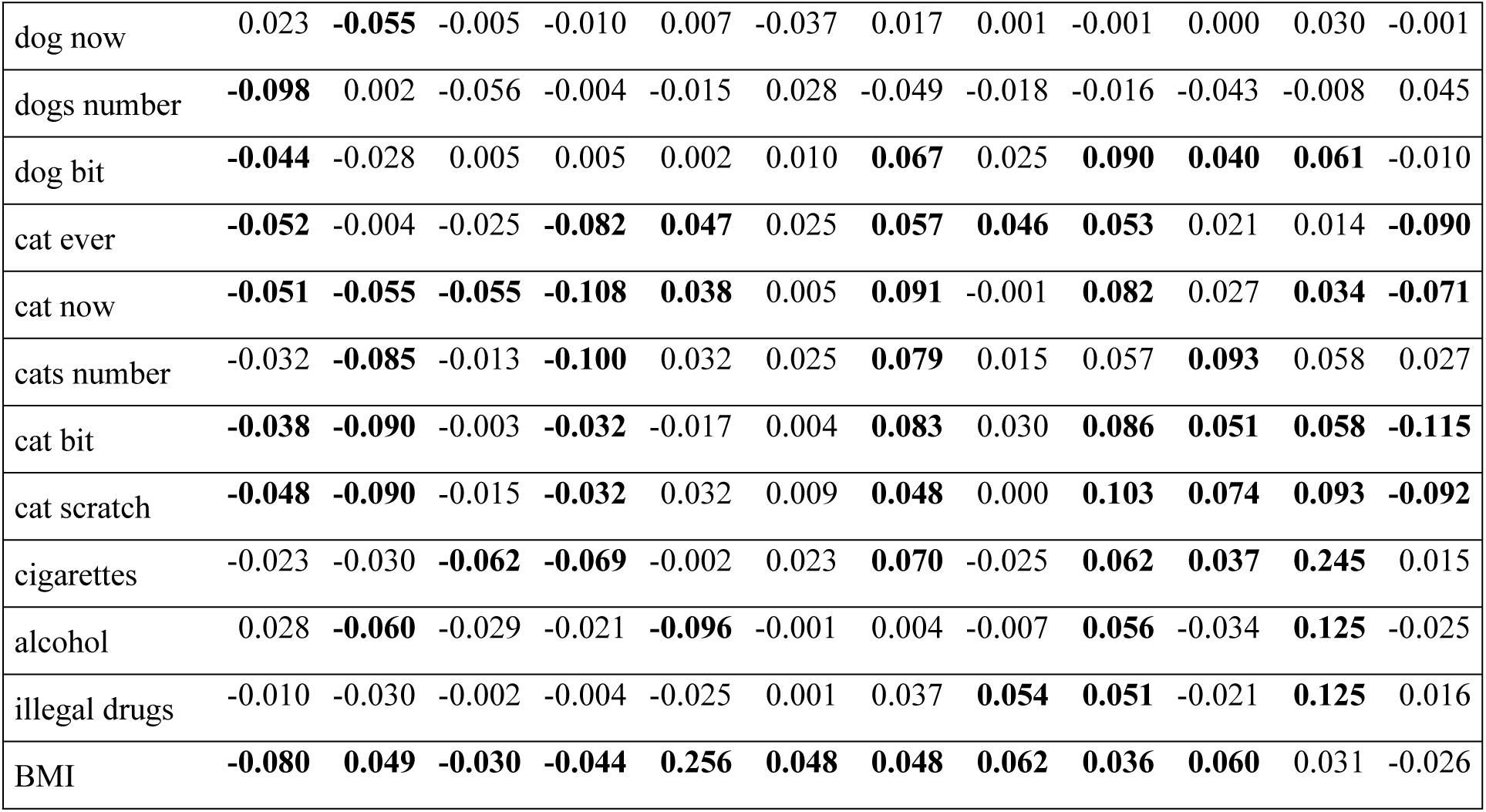
Association of animals-related variables with wellbeing and health of *Toxoplasma*-free female responders

## Discussion

Liking dogs, and especially liking dogs more than cats, was positively associated with some dimensions of quality of life as measured with WHOQOL-BREF, and with mental health in both men and women. This effect was stronger in older people. However, keeping dogs and cats, and even more sustaining animals-related injuries, correlated negatively with the quality of the responders’ life, with their physical health, mental health and biological fitness as measured by their number of children. Statistically, the effects of animal-related injuries were mostly relatively weak; however, their strength was comparable with the effects of four well-known risk factors, i.e., smoking, consuming alcohol, consuming illegal drugs, and high body mass index. In the present study, we also confirmed the existence of a strong association between latent infection with the parasitic protozoon *Toxoplasma gondii* and mental and physical health. However, we also showed that infection with *Toxoplasma* was not responsible for the observed association of wellbeing, health, and fitness with contacts with cats and dogs.

A statistically detected association between A and B does not mean causation. The negative correlation between seriousness of sustained cat-related injuries reported by the responders (A) and wellbeing score (B) could be the result of an association of both A and B with a third latent variable C, here, for example, the general pessimism of the responder. Even if the causality relation between A and B did exist, a cross-sectional study cannot discriminate between causes and effects. For example, a positive correlation between seriousness of sustained cat-related injuries reported by responders and number of diagnosed mental health disorders could be explained by a higher probability of cats to bite and scratch mentally ill people as well as by, for example, transmission of pathogens that are (co)responsible for mental illness, e.g. *Toxoplasma* [43] or *Bartonella* [44, 45], from cats to injured subjects. Regardless of these obvious methodological problems, even the data from observational studies could render some models/hypotheses more and some less feasible. For example, our data strongly contradict the model that transmission of *Toxoplasma* by a cat-related injury rather than the cat-related injuries themselves could be responsible for the negative association between cat-related injuries and mental health because this negative association was also detected in subjects who were *Toxoplasma*-seronegative. Theoretically, even this subset could contain a small fraction of subjects who acquired the infection after their serological testing for toxoplasmosis [46], and a larger but still rather small fraction of subjects who acquired the infection very long ago and their level of specific antibodies decreased under the arbitrarily set threshold of seropositivity [40]. If the presence of these false negative subjects was responsible for the observed association than the association should be much weaker in the seronegative subjects than in the whole population that contains not only the same fraction of false negative subjects, but also a large fraction of truly seropositive subjects. The opposite, however, was true – the strength of the observed effects (partial Kendall Tau reflecting the fraction of variability in mental health explained by the variables animals-related injuries) was much higher in *Toxoplasma*-free subjects than in the whole population. This suggest that both the pet-related injuries and toxoplasmosis have independent negative effects on mental health or mental health has independent effects on the probability of being injured by animals and being infected by *Toxoplasma* – see above. Similarly, the positive association between the number of cats in the home and the number diagnosed and non-diagnosed mental health disorders cold be explained both by the negative effects of cats on mental health as well as by a higher willingness of subjects who keep large number of cats to admit mental illness. The latter explanation is, however, contradicted by the fact that number of cats in the home correlated more strongly (in men in fact only) with the number of partner’s mental health disorders than with the number of the responder’s mental health disorders. It is indicative that this is not true for the cat-related injuries, which, in contrast to the number of cats in the home, affect only the responder and not his/her partner. Here, the association of injuries with the number of mental health disorders was much stronger than with the number of partner’s mental health disorders; in women, the latter association did not exist at all (Table S2).

The analyses performed on subjects who never kept a dog showed that, in men, a positive association between liking dogs and mental health and wellbeing (and negative with number of children, see below) could be the result of keeping a dog by people who like dogs. This conclusion, however, is only preliminary, as the positive association between liking dogs and wellbeing and with mental health still existed in this subpopulation but was much weaker than in the whole population (wellbeing Tau=0.011 vs 0.063; mental health Tau= −0.001 vs −0.026). On the contrary, the negative association of liking dogs with physical health, and also the negative association with number of children in women, also occurred in subjects who never kept a dog. Here, the situation was opposite than it was in men as these two associations were even stronger in women who never kept a dog than in the whole population (physical health: Tau 0.066 vs 0.021; children: Tau=-0.128 vs −0.095). No positive association between liking a dog and mental health was observed in women who never kept a dog (Tau=-0.001 vs 0.019). It seems that the positive effects of liking dogs in both men and women (namely better mental health) were related to a higher probability of keeping dogs by those who like dogs. On the other hand, the negative association between liking dogs and number of children in men and women as well as the negative association between liking dogs and physical health in women, have no direct relation to keeping a dog.

Different patterns showed in the analyses performed on subjects who never kept a cat. In men, a negative association between liking cats and mental health also existed in this subpopulation and was only slightly weaker than that observed in the whole population (Tau=0.039 vs 0.044). Unexpectedly, a positive association between liking cats and physical health was observed in this subpopulation (Physical health problems score Tau=-0.039), which was reflected by an even stronger positive association between preferring dogs to cats and physical health problems (Tau=0.054). These two associations were not detected in the whole population (liking cats Tau=-0.004; preferring dogs to cats Tau=0.008). In women, the strength of the association between liking cats and mental health problems was largely reduced (Tau=0.023 vs. 0.044) and with number of diagnosed and non-diagnosed mental disorders disappeared entirely (diagnosed: Tau=-0.003 vs. 0.048; non-diagnosed: Tau=0.015 vs. 0.045; total: Tau=0.007 vs. 0.055). It can therefore be suggested that not the liking of cats but the keeping of cats could be responsible for the association between liking cats and mental health problems in women and probably also in men. However, the negative association between liking cats and number of children was relatively strong in cat-free men (Tau=-0.044 vs. −0.082) and even stronger in cat-free women (Tau=-0.109 vs. −0.080), suggesting no direct relation to keeping a cat. It could be speculated that loving cats canalize parental instincts, and therefore negatively influence the fecundity of men and even more of women. Of course, opposite causation (strengthening of love for cats by the absence of children), could be also responsible for the observed negative association.

All analyses were also performed on subpopulations of people who had never been injured by a dog or by a cat. The observed negative association between number of dogs in the home and physical health problems was even stronger (Tau 0.121 vs 0.079) in the subset of men who were not injured by a dog than in the whole population. The same was true for the observed negative association between number of dogs in the home and wellbeing in women (Tau=-0.064 vs. −0.046). Again, a slightly different pattern was revealed in the analyses of subjects who had never been injured by a cat. In men, the negative association between keeping a cat and mental health was absent in the cat injury-free subpopulation (mental health problems score: Tau=0.015 vs. 0.044). On the other hand, the strong positive associations between keeping a cat or number of cats in the home and number of diagnosed and un-diagnosed mental health disorders in the responder’s partner was also present in the cat injury-free subpopulation (partner’s diagnosed: Tau=0.037 vs. 0.036; non-diagnosed: Tau=0.099 vs. 0.055, Total: Tau=0.080 vs 0.060). In women, the negative association between keeping a cat and mental health, including the associations with the numbers of diagnosed and un-diagnosed mental health disorders, were only slightly weaker in the cat injury-free subpopulation (mental health problems score: Tau=0.043 vs. 0.053; diagnosed: Tau=0.040 vs 0.053; non-diagnosed: Tau=0.038 vs. 0.049, Total: Tau=0.46 vs 0.060). On the other hand, the negative association between number of cats in the home and wellbeing and between number of cats in the home and mental health was absent in this subpopulation (wellbeing: Tau=-0.008 vs. −0.044; mental health problems score: Tau=0.008 vs. 0.033). It can by concluded that the dog- and cat-related injuries most probably played an important role in the mental health of participants in our study; however, keeping dogs and cats alone also has negative effects on their mental health. Keeping dogs (ever, currently, or number in the home) was positively associated with number of children in the dog injury-free men. This positive association was observed only between ever keeping a dog and number of children in the dog injury-free women. Number of children correlated positively only with ever having kept a cat and only in cat injury-free women; however, the strength of the negative association between number of children and number of cats in the home was the same in the cat injury-free women and whole population of women (Tau=-0.040). Again, animal-related injuries are probably not responsible for the observed association between keeping pets and fecundity.

The causality relation between the *Toxoplasma* infection and impaired mental health has been confirmed by prospective cohort study [47, 48] and experimental infections of laboratory animals. Such confirmation of the direction of causality between animal-related injuries and mental health is not currently available. However, according to three of nine Bradford Hill criteria of causation [49] (Specificity, Biological gradient, and Analogy), the animal-related injuries seems to be more probably the cause then the effect of impaired mental health: (1) As was already published, the spectra of dog biting-associated disorders and cat scratching-associated injuries do not intersect [50]. Therefore, the association probably does not result from a higher probability of reporting feelings of hurt or injustice, including sustained injury, by subjects with neuropsychiatric disorders (or generally by subjects in bad psychological conditions). (2) The strength of the effect positively correlated with the intensity of exposure to the suspected risk factor, here the intensity of sustained animal-related injuries. (3) The cats and dogs are vectors of some pathogens that are known to have negative impacts on human mental health [45, 51, 52]. In the present study we showed that *Toxoplasma* was not the cause of the observed associations between the injuries and impaired health of the responders [45]. However, the participants were not tested for, e.g., *Bartonella henselae*, the agent of cat-scratch disease. This disease probably has many effects on the physical and mental health of patients and on the incidence of some mental health disorders, most strongly on the risk of major depression [53–56]. Many other, both known and unknown, pathogens could be transmitted from animals to humans either by biting and scratching, or by other forms of close contact. Of course, the infection-based explanation of the association of contact with cats and dogs with mental health problems must be considered just as a working hypothesis that should be tested in future studies.

Liking dogs and cats was strongly negatively associated with number of children. Most probably, having (and liking) one’s own children has a negative effect on the relative intensity of liking pets. An alternative and more alarming explanation, i.e. the existence of a negative effect of liking pets on, e.g., parental instincts, should be tested in future cohort studies by searching for an association between liking pets and intensity of desire to have children in childless people. Also, we plan to repeat our study after five years to see whether young childless women, registered members of the Lab Bunnies community, who originally reported the more intensive love for cats would report fewer children in the follow-up study than those who originally reported less intensive love for cats.

The number of children was also lower in subjects who presently keep animals and even more so in subjects who were more seriously injured by animals. Here, the decrease of biological fitness could be the result of impaired physical and mental health of affected subjects. Despite the negative effects on the number of children, animal-keepers, and even more so subjects who were more seriously injured by dogs or cats, were more sexually active. This could be explained by the adoption of a “quick life strategy” by people in bad mental and physical health conditions. The slow life strategy, i.e., the preferential investment in quality of offspring, to carefully search for the best available sexual partners and to accumulation of resources, including education, before starting the process of reproduction, pays more to healthy subjects. For the subjects with lower life expectancy it pays to adopt the quick strategy, to have a sex soon and, especially in men, with as many partners as possible. In the affected (animal-keeping or animal-injured) men, high sexual activity was accompanied by high sexual desire. In contrast, in women, the affected subjects had lower sexual desire regardless of their higher sexual activity. This suggests that psychological mechanisms responsible for the transition from a slow to fast life strategy probably differ in men and women. The slow and fast life strategies were outside of the scope of present study; however, the existence of a highly significant positive correlation between both physical and mental health problems and sexual activity was detected in women, regardless of their contacts with dogs and cats.

A major limitation of the present study is its observational character. To discriminate whether, e.g., keeping cats has a negative effect on mental health or whether impaired mental health has positive effects on the probability of getting and keeping cats, an experimental approach is required. It is necessary, e.g., to provide a random sample of volunteers a cat (a dog) and after several years of keeping a cat (dog), compare their mental health with that of the unexposed controls. As far as we know, only one such study was published. It was performed on a sample of 48 hypertensive individuals and demonstrated the positive effect of keeping pets for six months [57].

Our results suggest that the more objective indices of health, namely the scores of physical and mental health computed on the basis of concrete parameters such as number of prescribed drugs consumed daily, number of diagnosed mental health disorders etc., better correlate with the animal-related variables than the responders’ subjective assessment of physical and mental health statuses. This suggests that real status, rather than its subjective assessment, is responsible for the observed association between the dog- and cat-related variables and reported health. This is in agreement with previously published observations showing that regardless of the pet keepers’ claim that the animals provide them many psychological and physiological benefits, the standard questionnaires show no evidence for this [20, 58]. It must be emphasized that the subjective perception and interpretation of one’s own situation and general mood probably influence the way the participants respond to seemingly objective questions, such as how often they visited a primary care doctor or how often they took antibiotics. Probably, the only questions that were objective and that really reflect the health status of the cat- and dog-keepers were those on the number of mental disorders of the participants’ partners. Based on these questions we can conclude that having a dog or a cat in house negatively correlates with mental health, which corresponds to the conclusion of the whole study.

Also, toxoplasmosis status was self-reported by the participants of the study. It was shown that the information on toxoplasmosis status perfectly corresponds to the information in our file for subjects who were tested for toxoplasmosis in our lab [40]. However, about 60% of male and 70% of female participants of the present study were tested elsewhere. It is possible that some subjects misreported whether they are *Toxoplasma* infected or not. Similarly, some responders who were *Toxoplasma*-negative during their serological test could have acquired the infection in the time between serological test and participation in the present study. It must be emphasized, however, that presence of misdiagnosed subjects in the population can result in a Type 2, not a Type 1 error – it can increase the risk of failure to detect existing effects but not the risk of detecting non-existing effects.

We asked the subjects whether a dog (a cat) had ever been kept in their family. However, most subjects had already two families, the family to which they were born and their own new family. It is possible that keeping pets in original and new family are associated differently with the subjects’ wellbeing and health. Any future study should discriminate between these two kinds of families.

It is widely believed that keeping cats and dogs has a positive impact on the subject’s quality of life despite most studies performed during the past 20 years showing the opposite – for an excellent review see [17]. It seems that this opinion could be the result of the subjective interpretation of quality of life of people who like dogs and cats and therefore also keep them in their house. When more objective information on quality of life is collected, the effects of keeping dogs and cats and especially of being injured by them, on quality of life is mostly negative [31]. Until now, most published studies examined the effects of contact with pets on wellbeing and health of patients or seniors. The number of studies performed on general nonclinical populations is much lower and those studying effects of keeping cats are even rarer. As far as we know, the effect of pet-related injuries on quality of life was studied only in our lab and only one other group reported the effect of being bitten by a cat on the risk of major depression [59] and another group reported the effect of being bitten by a cat prior to age 13 on symptoms of schizotypy in adulthood [60]. The major problem of the studies that claimed to find positive effects of contact with pets on various facets of quality of life was the autoselection of the participants. The studies were advertised as studies of the effect of pets on human life. It is rather probable that people who like pets, keep them and believe that their pets positively influence many aspects of their lives preferentially participate in such studies. People who like pets and believe that pets positively influence the quality of life of pet-keepers are also present in controls, however, here their frequency is lower than in the pet-keepers. This flaw occurs even in studies that explicitly claim the opposite. For example, Bennett et al. [8] wrote “We were careful not to indicate that we were particularly interested in pet owners,” however, the study had been advertised as “being concerned with investigating the daily activities, emotions and perceptions of older adults living independently, either alone or with other persons or pets, in the community.” To eliminate this source of bias, our study was advertised as a study on “mystical thinking, superstitions, prejudices, religion and relation between various environmental factors and health and wellbeing.” The pet-related questions were buried in many hundreds of other questions of the more than 80-minute questionnaire. It is probable that the relation to pets had only a small impact, if any, on how participants responded to our questions on quality of life and mental and physical health. Therefore, our study probably more objectively reflects the situation in general nonclinical population than most previously published studies. It is also rather probable that most students of the effects of pet-keeping on quality of life like dogs and cats and therefore *a priory* believe that keeping pets has a positive effect on quality of life [17, 31] (which was true also for us when we registered this study in our grant proposal). It is possible that most people are reluctant to publish results that contradict their expectations [17]. Similarly, many editors and referees are probably less willing to endorse the publication of manuscripts that contradict general opinion, and very often their personal beliefs too.

## Conclusions

From the point of view of statisticians, the effect sizes of the observed associations are small – the cat- and dog-related variables explained just a small percentage of total variability of the output variables. For example, the effect of cat scratching on mental health explained about 2.5% of total variability in the mental health problems score in the total population (but more than 10% of variability in the *Toxoplasma*-free subpopulation). It is important to remember, however, that these effects are of similar strength as those of other risk factors – smoking, alcohol, illegal drugs, and high BMI. Most importantly, more than 50% of households in developed countries keep a dog or a cat and this number is continuously rising. Therefore, the real impacts of formally weak effects of keeping of pets on public health and wellbeing could be enormous and growing.

## Acknowledgements

We thank Charlie Lotterman for the final revisions of our text.

## Supporting information

**S1 Fig. Effect of keeping a dog now on physical health of men and women of different age**

The boxes, spreads, upper numbers and lower numbers show standard errors, standard deviations, numbers of subjects in particular category and p-values of two-sided t-tests, respectively.

**S2 Fig. Effect of keeping a cat now on physical health of men and women of different age**

The boxes, spreads, upper numbers and lower numbers show standard errors, standard deviations, numbers of subjects in particular category and p-values of two-sided t-tests, respectively.

**S3 Fig. Effect of keeping a dog now on mental health of men and women of different age**

The boxes, spreads, upper numbers and lower numbers show standard errors, standard deviations, numbers of subjects in particular category and p-values of two-sided t-tests, respectively.

**S4 Fig. Effect of keeping a cat now on mental health of men and women of different age**

The boxes, spreads, upper numbers and lower numbers show standard errors, standard deviations, numbers of subjects in particular category and p-values of two-sided t-tests, respectively.

**S5 Fig. Effect of dog biting on wellbeing of men and women of different age**

The boxes, spreads, upper numbers and lower numbers show standard errors, standard deviations, numbers of subjects in particular category and p-values of two-sided t-tests, respectively.

**S6 Fig. Effect of cat biting on wellbeing of men and women of different age**

The boxes, spreads, upper numbers and lower numbers show standard errors, standard deviations, numbers of subjects in particular category and p-values of two-sided t-tests, respectively.

**S7 Fig. Effect of cat scratching on wellbeing of men and women of different age**

The boxes, spreads, upper numbers and lower numbers show standard errors, standard deviations, numbers of subjects in particular category and p-values of two-sided t-tests, respectively.

**S8 Fig. Effect of dog biting on physical health of men and women of different age**

The boxes, spreads, upper numbers and lower numbers show standard errors, standard deviations, numbers of subjects in particular category and p-values of two-sided t-tests, respectively.

**S9 Fig. Effect of cat biting on physical health of men and women of different age**

The boxes, spreads, upper numbers and lower numbers show standard errors, standard deviations, numbers of subjects in particular category and p-values of two-sided t-tests, respectively.

**S10 Fig. Effect of cat scratching on physical health of men and women of different age**

The boxes, spreads, upper numbers and lower numbers show standard errors, standard deviations, numbers of subjects in particular category and p-values of two-sided t-tests, respectively.

**S11 Fig. Effect of dog biting on mental health of men and women of different age**

The boxes, spreads, upper numbers and lower numbers show standard errors, standard deviations, numbers of subjects in particular category and p-values of two-sided t-tests, respectively.

**S12 Fig. Effect of cat biting on mental health of men and women of different age**

The boxes, spreads, upper numbers and lower numbers show standard errors, standard deviations, numbers of subjects in particular category and p-values of two-sided t-tests, respectively.

**S13 Fig. Effect of cat scratching on mental health of men and women of different age**

The boxes, spreads, upper numbers and lower numbers show standard errors, standard deviations, numbers of subjects in particular category and p-values of two-sided t-tests, respectively.

**S14 Fig. Association between intensity of sustained dog biting and wellbeing – the scores of WHOQOL-BREF domains**

The categories on x-axis describing the intensity of being injured by a pet are: 0-never, 1-only while playing, 2-only as a warning, 3-yes, minor injury (only skin cut), 4-yes, moderate injury (bleeding), 5-yes, serious injury, I had to seek medical treatment.

**S15 Fig. Association between intensity of sustained cat scratching and wellbeing – the scores of WHOQOL-BREF domains**

The categories on x-axis describing the intensity of being injured by a pet are: 0-never, 1-only while playing, 2-only as a warning, 3-yes, minor injury (only skin cut), 4-yes, moderate injury (bleeding), 5-yes, serious injury, I had to seek medical treatment.

**S1 Table Distribution of responses on particular questions of the questionnaire – categorical and ordinal variables**

*The column 3 shows mean answer to particular questions and the columns 4-12 the percentage of subjects who provided particular answer (0-8). The questions on variables printed bold were responded by a code of the answer (for meaning of particular codes see the Material and Methods), other questions by a number (e.g. 4 children). In such cases, the highest number of the scale (e.g. 8) always means “eight or more”. The codes in parenthesis in the first column indicates whether particular variable correlate significantly with sex (s) age (a), education (e), and urbanization (u) in the whole population, men, and women, respectively. The code 0 means no correlation with sex, age education or urbanization*.

**S2 Table Continuous depending variables - difference between men and women**

*For the explanation of codes in the column 1 see the table 2*.

**S3 Table Effects of pets on health and wellbeing – Men**

**S4 Table Effects of pets on health and wellbeing – Women**

**S5 Table Effects of pets on health and wellbeing – Men who never kept a dog**

**S6 Table Effects of pets on health and wellbeing – Women who never kept a dog**

**S7 Table Effects of pets on health and wellbeing – Men who never kept a cat**

**S8 Table Effects of pets on health and wellbeing – Women who never kept a cat**

**S9 Table Effects of pets on health and wellbeing – Men who were never injured by a dog**

**S10 Table Effects of pets on health and wellbeing – Women who were never injured by a dog**

**S11 Table Effects of pets on health and wellbeing – Men who were never injured by a cat**

**S12 Table Effects of pets on health and wellbeing – Women who were never injured by a cat**

**S13 Table Effects of pets on health and wellbeing – *Toxoplasma*-free men**

**S14 Table Effects of pets on health and wellbeing – *Toxoplasma*-free women**

